# lncRNA MANCR isoforms selectively mediate multiple levels of epigenomic and P53-responsive transcriptional control in triple negative breast cancer

**DOI:** 10.64898/2026.04.06.716674

**Authors:** E. Pacht, J.S. Warren, R.H. Toor, K.C. Glass, H.W. Greenyer, A.J. Fritz, B. Banerjee, S.E. Frietze, J.B. Lian, J.A.R. Gordon, G.S. Stein, J.L. Stein

**Affiliations:** Department of Biochemistry, University of Vermont; Department of Pharmacology, University of Vermont; University of Vermont Cancer Center, University of Vermont

**Keywords:** long noncoding RNA, P53, genomic instability, microRNA (miRNA), breast cancer, epigenetics, transcriptional isoforms, MANCR, genomic instability, triple negative breast cancer

## Abstract

Long noncoding RNAs (lncRNAs) are important regulators of gene expression and are frequently dysregulated in cancer. The mitotically associated lncRNA MANCR is highly expressed in aggressive cancers and contributes to genomic instability in triple-negative breast cancer (TNBC), but the molecular mechanisms underlying its activity remain poorly defined. Here we integrate computational and experimental approaches to examine the structure and regulatory interactions of MANCR isoforms. Analysis of transcriptomic datasets revealed tumor-type-specific expression patterns for seven MANCR isoforms in breast cancer cell lines. Computational prediction of RNA secondary structures identified conserved structural features across isoforms, suggesting potential functional specialization. We identify p53 as a MANCR-interacting protein through computational docking and RNA immunoprecipitation sequencing (RIP-seq) and demonstrate that MANCR depletion reduces p53-dependent transcriptional activity. Chromatin isolation by RNA purification sequencing (ChIRP-seq) revealed 1, 250 genomic regions associated with MANCR, including enrichment of p53 consensus motifs and GC-rich sequence elements. Motif analysis further identified candidate sequence features associated with MANCR-occupied chromatin regions. Computational prediction of RNA-miRNA interactions identified multiple potential miRNA binding sites across MANCR isoforms, including miR-6756-5p, which targets the androgen receptor (AR). Consistent with this prediction, AR expression decreased following MANCR knockdown in TNBC cells. Together, these results suggest that MANCR isoforms may contribute to transcriptional regulation in TNBC through interactions with chromatin, p53 signaling pathways, and potential miRNA regulatory networks.

**One Sentence Summary:** Mitotically-associated lncRNA (MANCR) is prevalent in aggressive cancers interacting with DNA, P53, and miRNAs, to mediate multiple levels of epigenetic transcriptional control in triple negative breast cancer.

## Introduction

Long-noncoding RNAs (lncRNAs) are key to biological regulation of critical cellular pathways, including proliferation, apoptosis, invasion, and cell migration (1). Major regulatory roles of ncRNAs have been implicated in diverse diseases including cancer, Alzheimer’s disease, and Down syndrome (2–4). In addition to independent roles of lncRNAs, they can form complexes with DNA, protein, and RNAs (5). These complexes include networks of chromatin-regulating enzymes and splicing factors (6). LncRNAs induced by hypoxia have been shown to promote alternative splicing, functionally linking lncRNAs to hypoxia-induced alternative splicing in tumorigenesis (7, 8). Long ncRNAs (lncRNAs) adopt defined structures that contribute to biological regulation of gene expression and are implicated in cancer-compromised transcriptional and epigenetic control (7, 9–11).

Dysregulation of lncRNAs has been associated with oncogenesis and tumor progression (12). Mitotically associated lncRNA (MANCR) is a lncRNA associated with aggressive breast cancer (13). In triple negative breast cancer (TNBC) cells, depletion of MANCR significantly reduces cell viability as a consequence of increased DNA damage. MANCR was discovered due to elevated expression and chromosome association in mitotic cells, although other studies have shown that it modulates cell cycle progression through regulation of RhoGEF (14). MANCR expression in metastatic TNBC is further increased following cellular stress and under hypoxic conditions (14). MANCR has been shown to interact with all human autosomes and the X chromosome, which is consistent with MANCR playing a unique role in maintaining genomic stability during cancer progression (13). Depletion of MANCR is associated with reduced lung metastasis in vivo, and decreased cell migration, invasion, wound healing, defective cytokinesis and increased cell death, reinforcing engagement in stabilizing the cancer cell genome (14). It is clear that MANCR has diverse roles regulating multiple cellular functions. Among cancer-associated lncRNAs, alternative transcriptional start and end sites have been associated with altered lncRNA isoform structures and specific functions in cancer initiation and progression (15). A compelling mechanism contributing to the diverse functionality of MANCR could reside in isoform expression as well as the secondary and tertiary structure of the specific isoforms.

Developing an understanding for the structure of lncRNAs can contribute to functional characterization. This expectation is supported by structural analysis of other long-noncoding RNAs, which include Metastasis-Associated Lung Adenocarcinoma Transcript 1 (MALAT1), a nuclear-localized lncRNA with complex oncogenic roles (11). Structural studies identified pseudoknots, structured tetraloops, and structured internal loops as well as intramolecular long-range interactions in these RNAs (11). Cancer-associated mutations are predicted to alter the structure of other lncRNAs and expose binding sites for microRNAs associated with cancer (11). The dynamic structure of lncRNAs can inform the functions of cancer-associated mutations through altered microRNA interactions (11). Further computational approaches, combined with FRET and NMR spectroscopy, identified unique binding of multiple small molecules to MALAT1 that induced structural change of the lncRNA and subsequently decreased non-coding RNA levels are accompanied by changes in morphogenesis in an organoid model (16). Specificity is indicated by the absence of effects on NEAT1 (17). Recently, another lncRNA has been shown to bind to RNPS1, colocalize to nuclear speckles, and regulate alternative splicing (18). Computational observations identified stable interactions between MALAT1 and RNPS1 in a loop-specific manner, establishing the potential to target this complex for therapeutic application (18).

Recently, several lncRNAs have been demonstrated to interact with P53 in cancer models to regulate signaling and downstream effects associated with P53 function (19–21). Mutant P53 has been shown to deregulate genes associated with Epithelial-Mesenchymal Transition (EMT) and chemoresistance in breast cancer (22). Complexes with lncRNAs and miRNAs have been identified as components of feedback loops for P53 and have been suggested as prognostic indicators (23, 24). The lncRNAs associated with these networks varied in expression by tumor sub-type and included lncRNAs known to be associated with TNBC that include DINO, MALAT1, and PANDA (20, 21, 25–28). Basal tumors with mutant P53 contained downregulated Androgen Receptor (AR) and exhibited higher EMT scores, associated with poorer prognosis (22). Expression of AR was rescued through overexpression of wildtype P53 that led to decreased expression of mesenchymal markers. The association between AR and P53, and the lncRNA-mRNA-miRNA networks may play a role in EMT and chemoresistance in breast cancer (22). Another mechanism for lncRNA activity is preventing cell senescence through the degradation of P53 by LINC02593 (29). This study demonstrates the ability of lncRNA LINC02593 to scaffold interactions between the coiled-coil domain of COP1 and the P53 C-terminus, highlighting a mechanism for lncRNAs to bridge proteins leading to cell survival and continued cancer growth (29).

Although modulation of MANCR expression and activity has been shown to have a profound effect on gene expression and cell survival, the mechanism by which MANCR modulates these key parameters of biological control are unresolved. Here we report predictions of MANCR structure that propose MANCR-mediated mechanistic control of genomic stability and cell survival in breast cancer cells. Computationally identified structures are experimentally validated by demonstrating MANCR-mediated regulation of gene expression, engagement in DNA and protein interactions and control of microRNA stability that impacts P53 regulatory pathways.

## Results

### Selective MANCR isoform expression in breast cancer cell lines

High MANCR expression is associated with aggressive and metastatic cancers with worse prognosis and lower survival, with a hazard ratio of 2.489 (p-value = 1.29e-7) (30) (Fig 1A). It has previously been shown that MANCR has a role in TNBC (13). Numerous cancer cell lines have high MANCR expression, however a frequently used TNBC cell model MDA-MB-231 exhibits the highest MANCR expression reported for TNBC (Fig 1B). MANCR has been reported to have multiple transcript isoforms with different initiation sites and gene lengths (Fig 1C). Other studies have shown that lncRNA isoforms exhibit unique functional cancer-related roles (15). GENCODE Release 45 (GRCh38) identified seven major MANCR transcripts (31). The seven isoforms we examine differ substantially in length. MANCR isoform 201 (1, 528 bp) is the longest isoform, followed by 204 (1, 448 bp) and 206 (1, 297 bp). Intermediate-length MANCR isoforms include 203 (703 bp), 205 (615 bp), and 207 (450 bp), whereas the shortest MANCR isoform is 202 (418 bp). The variety of isoform length supports potential structural and functional diversity. We evaluated the abundance of MANCR isoforms in 11 breast cancer cell lines and observed variations in isoform-specific distribution patterns (Fig 1D). In breast cancer cell lines the MANCR-201 isoform was most abundant in MDA-MB-157, Hs578T and BT-549, which are all TNBC cell lines. Similarly, MANCR-201 is the most abundant isoform in MDA-MB-231 cells (Fig 1E). In contrast, MANCR-205 was predominant in MDA-MB-436 and MDA-MB-468 TNBC cell lines (Fig 1D).

**Fig. 1.**
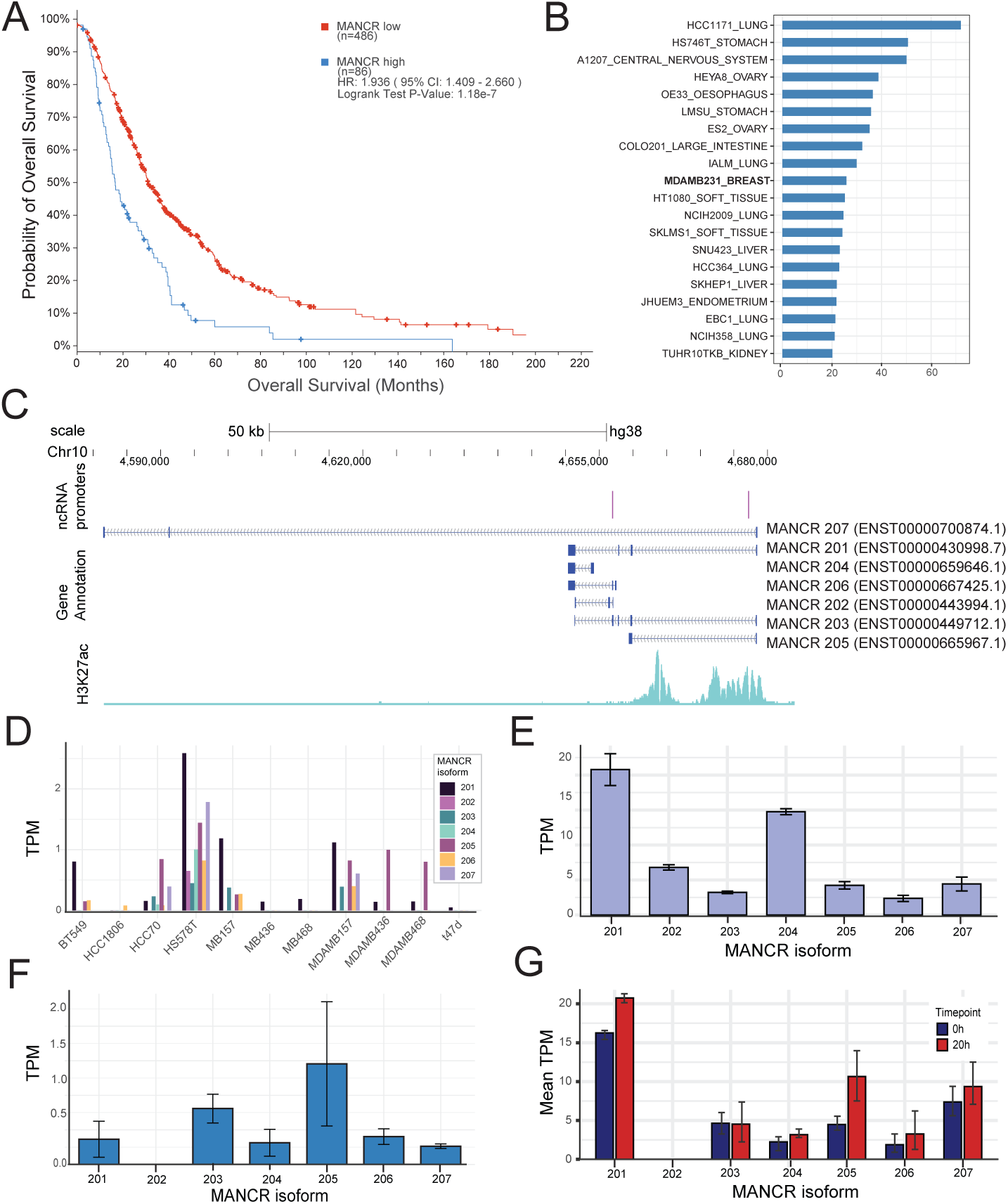
MANCR is expressed across cancer types and isoform expression is influenced by condition and cell line. (A) Kaplan-Meier curve of survival of pan-cancer patients with highly aggressive tumors stratified by MANCR expression 84 patients of high MANCR expression (RSEM > 0.5) and 84 patients with low MANCR expression. Statistical significance was determined using logrank test (p = 1.62e-7) (B) MANCR RNA expression across cancer cell lines from Cancer Cell Line Encyclopedia (CCLE). (C) MANCR isoform expression in breast cancer cell lines. Values are expressed as TPM. (D) MANCR RNA isoform expression in MDA-MB-231 cells. (E) MANCR isoform expression in MCF10A cells following the degron-mediated loss of RUNX1. Error bars represent SD from three replicates. (F) MANCR RNA isoform expression following antimetastatic drug NAMI-A treatment in MDA-MB-231 cells. Data represents mean of two replicates (47).

Expression of MANCR isoforms can be selectively altered by culture conditions and treatments. Normal MCF10A mammary epithelial cells do not express MANCR isoforms at detectable levels. However, in normal mammary epithelial cells undergoing EMT induced by removal of the RUNX1 tumor suppressor (32) MANCR expression is concurrently upregulated, and we identify 4-fold higher expression of MANCR-205 compared to -201 (Fig 1F). This result suggests that altered MANCR isoform expression accompanies different mechanisms of oncogenic transformation. Supporting this isoform switching, altered MANCR isoform expression is observed in MDA-MB-436, MDA-MB-157, and HCC70 cell lines under hypoxic conditions (Supp Fig 1) (33). Following hypoxia treatment, MANCR-205 was the predominant isoform in each cell line, even when it was not the predominant isoform prior to treatment (Supp Fig 1). NCBI Sequence Read Archive associated with BioProject PRJNA268434 was obtained to assess MANCR expression associated with NAMI-A metastatic drug in triple negative breast cancer. In MDA-MB-231 cells, treatment with the antimetastatic cancer drug NAMI-A increases MANCR expression, with MANCR-205 increasing approximately 2.3x (Fig 1G). The increase in MANCR-205 is associated with several parameters of cancer progression, including EMT, response to hypoxia, and drug resistance. Altered expression of MANCR isoforms highlights potential isoform-specific functions and the propensity for specific roles of MANCR isoforms across cell lines.

### MANCR isoform 2D structures

The function of lncRNA can be informed by 2D structures (11, 15, 16). To gain new insights into the function of MANCR isoforms we used RNAfold (34) to predict their 2D structures (Fig 2A-B, Supp. Fig. 2). The 2D structure predictions for MANCR isoforms identifies two novel structures for MANCR-201 and -205, with the base pairing probability color coded for the minimum free energy conformations (MFE). (Fig 2A-B). Based on the MFE models, it was observed that MANCR-201 and -205 exhibit unique structural features, including an Apex, Trunk, Core (containing 3’ and 5’ ends), and three distal protruding Arms. MANCR-201 formed a highly stable structure (MFE=-419.20), whereas MANCR-205 had a less stable structure with a higher free energy conformation (MFE=-186.60). However, MANCR-205 contained overall, higher base pairing probabilities than MANCR-201 (Fig 2A-B, Table 1). Regions with the highest mean base pairing probabilities in MANCR-201 include Arm 1 (0.865), and Arm 2 (0.884) (Fig 2A, Table 1). In MANCR-205, the highest mean base pairing probability was the Trunk (0.931) and the Core (0.861), while Arm 1 had a high mean base pairing probability (0.842), suggesting these are the most stable regions of this isoform (Fig 2B, Table 1).

**Fig. 2.**
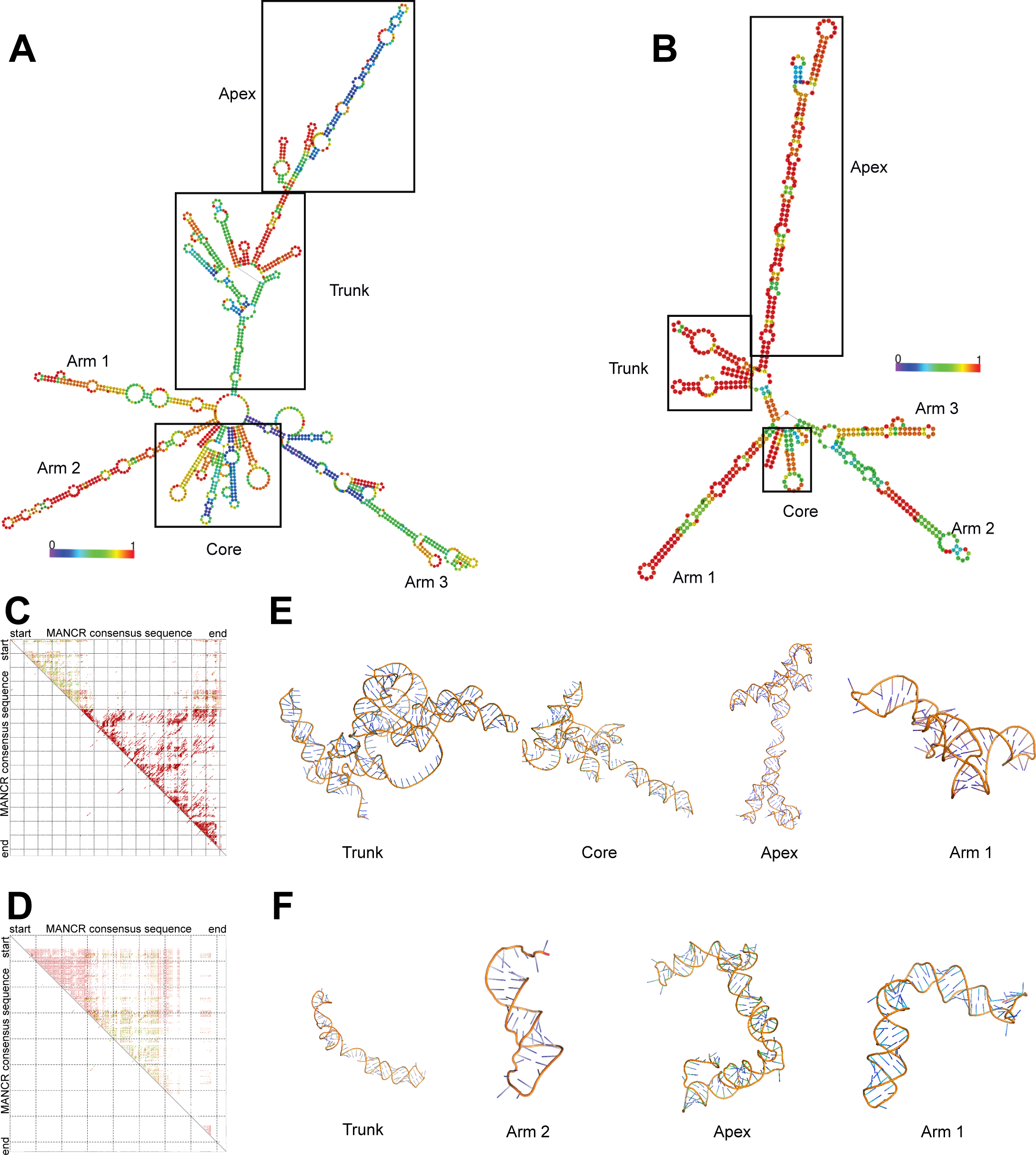
Structural prediction of MANCR isoforms. Base pairing probability of RNAfold 2D predicted structures for MANCR isoforms MANCR-201 (A) and MANCR-205 (B). Color scale represents base pairing probability for minimum free energy (MFE) structure with red indicating high base pairing probability and blue indicating low base pairing probability. Identified regions based on structural stability are highlighted (boxed) and labeled. Conservation of structural stability across MANCR isoforms. RNAfold 2D prediction dot plots on consensus sequences for MANCR-201 -204 -206 (C) and MANCR-202 -203 -205 (D) based on MFE. Plots indicate a combination of values from structure prediction, comparative sequence alignment, base pairing frequency, and pseudo entropy. Red indicates one type of base pairing between isoforms, ochre represents two types, and green represents three types of base pairing between isoforms. Base pairs compatible with the highest ranked pairs are shown on the lower left side of the plot. 3D structure predictions of highly stable regions were generated by RNAComposer for MANCR-201 (E) and MANCR-205 (F) based on base pairing probabilities and MFE. High stability regions were defined as having over 90% base pairing probability and also shared consensus structure stability.

Homology modeling indicated that MANCR-201, -204, and -206 share sufficient sequence similarity to generate a shared consensus structure (Mean pairwise identity: 84.84%, SCI=0.703, MFE=-266.28) (Fig 2C). Similarly, MANCR-202, -203, and -205 share sufficient sequence similarity to develop a consensus structure (Mean pairwise identity: 55.60%, SCI=0.141, MFE -23.68) (Fig 2D). The isoform MANCR-207 sequence and predicted 2D structure are unique, suggesting that MANCR-207 may be structurally unique from the other MANCR isoforms. Expression of MANCR-207 was consistently low across cell lines, and it was therefore excluded from further structural analysis. In the MANCR-201 consensus structure (which includes MANCR-204 and -206), sequence conservation was observed in the Core (bases 627-889), Arm 1 (bases 270-365), Trunk (bases 9-270 and 1103-1217), and the Apex (bases 1217-1497) (Conservation >90%). In MANCR-205 consensus structure (which includes MANCR-202 and -203), the regions with sequence conservation above 90% were limited to the Trunk and lower portion of the Apex (bases 343-486). Identified domains of structural conservation (stems) were identified with the RNAalidot program (34) (Fig 2C-D). In the MANCR-201 consensus structure, a total of 91 stems with conserved base pairing with an average probability above 0.5 were identified (Supp Table 1). Arm 3 and the Core contained the highest number of stems (40 and 22 respectively), although these stems were short with an average base pairing of 2.4 and 2.9 and mean probability scores 0.676 and 0.62.5 respectively. This structural landscape suggests that although the length of stems is relatively short, the high number of conserved stems and structural conservation across these regions establish that MANCR-201, -204 and -206 share significant structural stability. Also supporting this conserved stability, the Trunk region contains 12 conserved stems with a mean base pairing probability of 0.725. In contrast, we identified only 15 conserved stems across MANCR-205, -202, and -203 with a probability threshold ≥ 0.5 (Supp Table 1). The Apex contained the highest number of stems (8) with a mean probability of 0.60, whereas the Trunk contained only two conserved stems with a high average probability of 0.891. Although the number of conserved base pair stems and pairing probabilities of the MANCR-205 consensus structure is largely lower than the MANCR-201 consensus structure, both of these structures are likely to be the major predicted stable conformations of the MANCR isoforms.

As MANCR-201 and -205 are the major contributors to the identified 2D consensus structures, these isoforms were used for 3D modeling to identify 3D packing of structures. The core and trunk domains of MANCR-201 and -205 displayed 3D packing of hairpin structures (Fig 2E-F). In MANCR-201, the Apex domain was predicted to have a single strand region available for base pairing, consistent with a flexible conformation and supported by low base pairing probabilities in the 2D structure. Although MANCR-201 and -205 have substantially different predicted 2D and 3D stability and domain packing, it is clear the both isoforms (and by extension MANCR-204, -206, -202, -203) have conformationally stable domains that would suggest different functionality for MANCR isoforms that could include unique protein and nucleic acid interactions that are domain specific.

### MANCR P53 interactions directly regulate transcription

Based on our identification of MANCR domains that have a high probability of interacting with proteins and reports that other lncRNAs have been shown to interact with P53, including DINO, MALAT1, PANDA, we examined the ability of MANCR to interact with P53 (20, 21, 25–28). Interactions of MANCR with P53 were explored based on 3D structures of MANCR that are consistent with functional implications for interactions of P53.

Full-length MANCR isoforms and individual stable and unstable regions of MANCR-201 and MANCR-205 were computationally interrogated for P53 binding by AlphaFold (35). Although AlphaFold could not confidently identify a stable structure for the full-length transcripts, when MANCR-P53 interactions were predicted by individual MANCR domains the program identified the Arm 2 region of MANCR-205 as a potential P53 binding region. This is consistent with our finding that the Arm 2 region of MANCR-205 has a unique structure that is not predicted to bind DNA (Fig 6C), suggesting a specific protein binding function for this region. Although AlphaFold is sufficient to identify protein-protein interactions, the computational requirements of RNA folding required analysis by the HDOCK algorithm (36). Using known P53 structures in the Protein Data Bank with PDBIDs 2OCJ (Supp. Fig. 3A) and 2ADY (Fig 3A) we modeled docking of Arm 2 of MANCR-205 to determine if there were a stable interaction both in the presence and absence of DNA. The HDOCK analysis demonstrated a significant interaction with P53 in the presence of DNA with a high confidence score (0.9861) (Fig 3A). Intermolecular interactions were defined as distances under 3 Å between MANCR and P53 residues. There was substantial similarity in which MANCR bases interacted with P53 with and without DNA present, and the distal region of MANCR Arm 2 was identified as the interacting region. Specifically, MANCR bases 69-72 and 83-90 displayed close interactions with P53, with bases 69-72 appearing to directly interact with DNA as well. The P53 residues corresponding to the MANCR binding region are amino acids 121-184. This finding suggests that MANCR has a role in stabilizing P53-DNA interactions and may play a role in mediating gene expression.

**Fig. 3.**
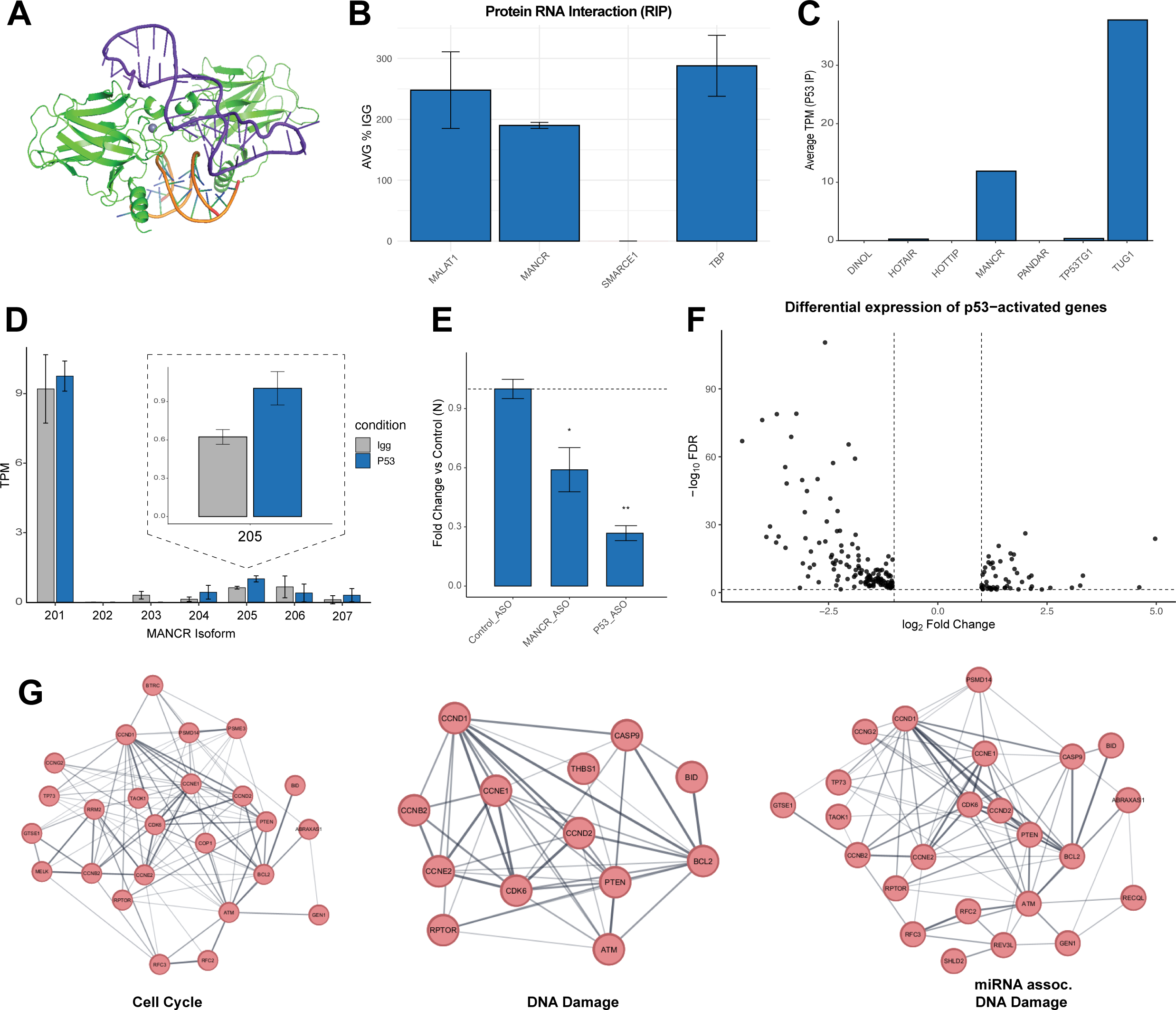
MANCR is predicted to interact with P53. (A) 3D modeling of P53 interactions with MANCR isoform 205. The P53 dimer structure (PDBID: 2ADY) is represented in green, with metal ions in gray and DNA in yellow. Arm 2 of MANCR-205 is represented in purple. Docking was performed with HDOCK and visualized in PyMol. (B) MANCR is enriched over IgG in P53 RIP-qpcr. MALAT1, MANCR, and TBP all showed significant enrichment with FDR < 0.05. (C) P53-associated lncRNA TPMs from P53 RIP-seq in MDA-MB-231 cells. (D) MANCR-205 is enriched over IgG in P53 RIP-seq. (E) P53 transcriptional activity decreases following knockdown by gapmers of MANCR and P53, shown by luciferase assay. (F) Differentially expressed genes that are known to be activated by P53 identified by RNA-seq following MANCR knockdown tend to be downregulated. (G) Pathway visualization of differentially expressed genes known to be P53 activated.

To experimentally confirm the interaction between MANCR and P53, RNA Immunoprecipitation (RIP)-qPCR was performed in MDA-MB-231 cells. Following immunoprecipitation of P53 all interacting RNAs were isolated and MANCR was identified as an interacting RNA with a significant enrichment over IgG (FDR=0.004, FC=1.88) (Figure 3B). Additionally, significant enrichment over IgG of MALAT1 and TBP were also identified, consistent with previously reported results showing that MALAT1 and TBP interact with P53 (25, 37). Exon structure of the MANCR gene does not allow for confident isoform-specific analysis by qPCR, therefore we identified isoform-specific P53 binding by unbiased RIP-seq in MDA-MB-231 cells. Quantification of total RNA associated with P53 was calculated as transcripts per million (TPM) (Fig. 3C) and identified 797 RNAs enriched over IgG (Log2FC > 0.5, p-value < 0.05). This analysis confirmed that MANCR was substantially associated with P53 (TPM=11.9). Additionally, lncRNAs previously demonstrated to interact with P53 (eg TUG1, DINO, HOTAIR) were identified in this analysis (Fig 3C) (20, 38, 39). Of the specific MANCR isoforms, MANCR-201 was the most abundant, followed by MANCR-205, although MANCR-201 showed non-specific interaction with IgG whereas MANCR-205 was preferentially enriched over IgG (Fig 3D-E). Reinforcing this finding, publicly available data from NCBI Sequence Read Archive associated with BioProject PRJNA1210463 demonstrates that P53 interacting RNAs in HCT166 colorectal cancer cells identified lncRNA LFPM interactions with P53 by RIP-seq. In this study a specific LFPM loop structure that directly interacts with the DNA-binding domain of P53R175H to stabilize P53 activity, thus protecting cell proliferation and resistance to ferroptosis was reported. Our analysis of this dataset identified that MANCR was associated with both wild type and R175H mutant P53 (Supp. Fig. 3B). Similar to our findings, MANCR-201 and MANCR-205 were strongly associated with P53. The most abundant isoform interacting with wild type P53 was MANCR-201 (1.04 TPM), followed by MANCR-205 (0.87 TPM), similar to patterns we identified in MDA-MB-2321. MANCR-205 showed substantial variation in transcript abundance (TPM) between wild type and mutant P53 (0.47 TPM) that does not occur with other isoforms suggesting differential affinity of binding with specific P53 mutants with MANCR-205 (Supp. Fig. 3B). Notably, the R175H p53 mutation occurs within the predicted MANCR-205 binding region, further supporting the biological relevance of this interaction.

We directly address the effect of MANCR-P53 interactions on transcriptional activity by a P53 consensus-motif driven luciferase reporter construct. MDA-MB-231 cells containing a P53 luciferase reporter were treated with a cohort of gapmers to comprehensively target pan-MANCR isoforms or P53 or negative control (ASO). A significant reduction (p-value = 0.0022) in P53-mediated luciferase activity was observed following knockdown of P53 compared to negative control (0.27 fold change), corresponding to a 73% reduction in P53 activity (Fig 3F). Knockdown of MANCR similarly resulted in a significant (p-value = 0.018) reduction in P53-mediated luciferase activity (0.59 fold change), which corresponds to a 41% reduction in P53 transcriptional activity compared to negative control (Fig 3F). These results suggest that MANCR is directly involved in P53-mediated transcriptional activity.

To directly identify endogenous P53-regulated genes we performed RNA-seq in MDA-MB-231 cells prior to and following MANCR knockdown. We examined genes known to be activated by P53 (40) in the presence or absence of MANCR. There was an overall decrease (Log2FC <-2, FDR < 0.05) in P53-activated gene expression (130 of 185 genes) following MANCR loss (Fig 3G). P53-regulated genes affected by MANCR knockdown are involved in DNA damage responsiveness (eg. PTEN, RPTOR, and TP73), cell cycle control (eg. CCND2, CDK6, and CCNE1), and miRNA-associated DNA damage (eg. ATM) all of which are fundamental mechanisms driving cancer initiation and progression (Fig 3H).

The association between MANCR and P53 alterations has clinical correlates as amplification of MANCR is observed among cancers with high P53 mutational burden. These cancers include breast, lung, bladder and ovarian tumors and highly aggressive and metastatic tumors (Fig 4A-B) (30). In breast cancer, statistical analysis to determine if MANCR amplification and P53 mutation co-occur or are mutually exclusive demonstrated that MANCR alteration is significantly associated with P53 mutations (p-value <0.001, Log2 Odds Ratio 2.224). Specifically, MANCR amplification co-occurs with P53 missense mutations and P53 truncations in breast tumors (Fig 4C). The alteration frequency of P53 in breast cancer patients with high MANCR expression is nearly 70%, substantially higher than the alteration frequency in patients with low MANCR expression (Fig 4D). Further, the alteration frequency of P53 in patients with high MANCR expression was notably higher than comparable genes (Fig 4D). These data would strongly suggest a functional link between MANCR expression and P53 in breast cancer patients.

**Fig. 4.**
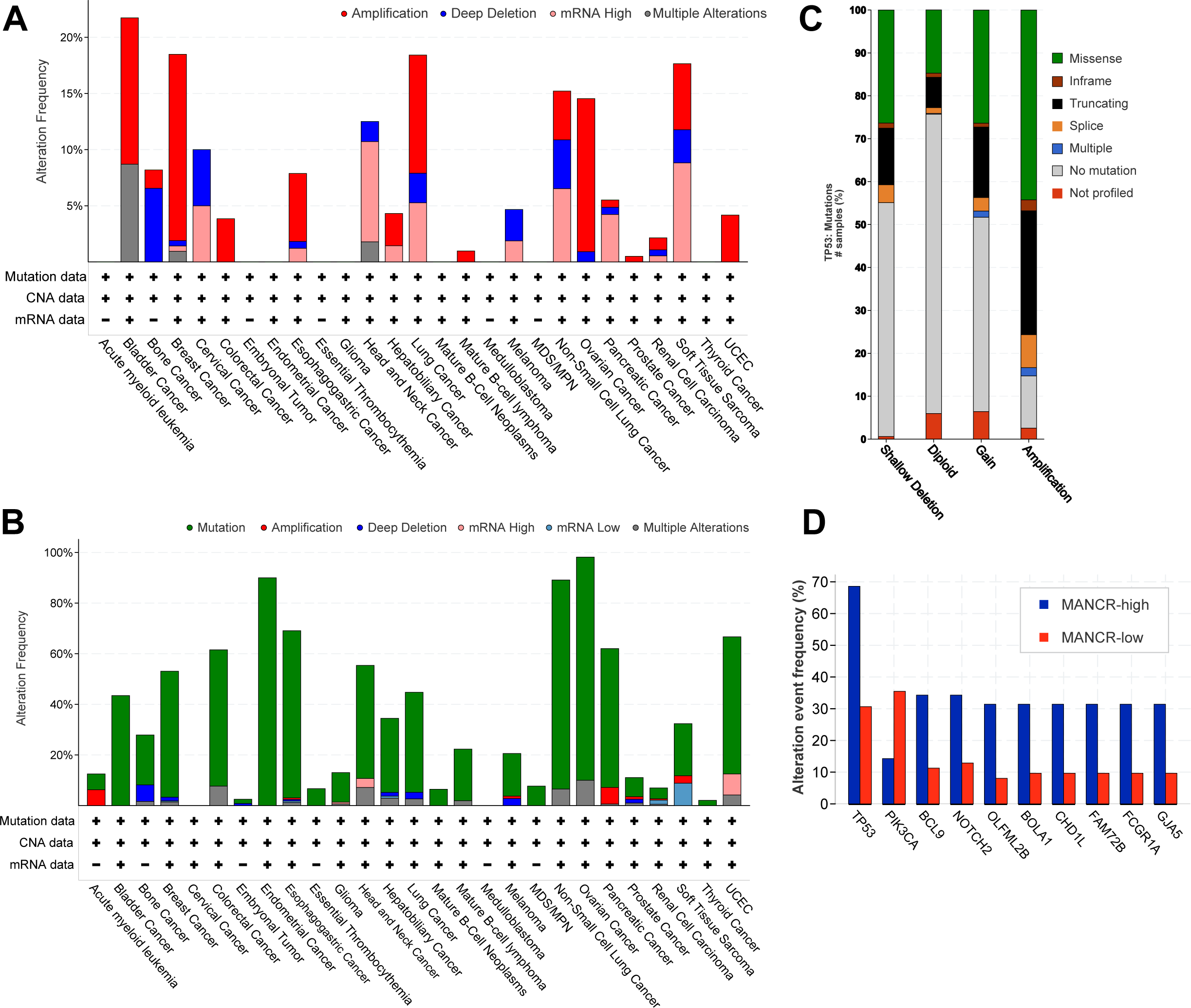
MANCR amplification and P53 mutation are co-occur in patients. (A) CBioPortal bar graphs of MANCR alterations across TCGA pan-cancer patients (B) CBioPortal bar graphs of P53 mutations across TCGA pan-cancer patients. (C) CBioPortal bar graph of P53 mutations by MANCR alterations within breast cancer patients. (D) Alteration frequency of genes associated with MANCR in invasive breast cancer.

### MANCR directly interacts with genomic targets

Interactions between MANCR and P53, and previous findings that MANCR is chromosome associated (13), suggest an epigenomic role for this lncRNA. Therefore, we performed ChIRP-seq to directly characterize MANCR binding across the genome. We identified 1250 MANCR binding sites throughout the genome (Fig 5A). A large subset of MANCR binding regions were found to be associated with P53 consensus sequence by HOMER motif analysis (q-value < 0.0001, 5.1-fold enriched over background) (41). The P53 consensus sequence was identified as the most significant (q-value < 0.0001) transcription factor binding motif in MANCR ChIRP-seq. Enrichment of p53 binding motifs identified in MANCR ChIRP analysis were confirmed by FIMO motif analysis (42), which substantiated 116 P53-motif occurrences with high significance (p-value < 0.0001) (43) (Supp Table 2).

**Fig. 5.**
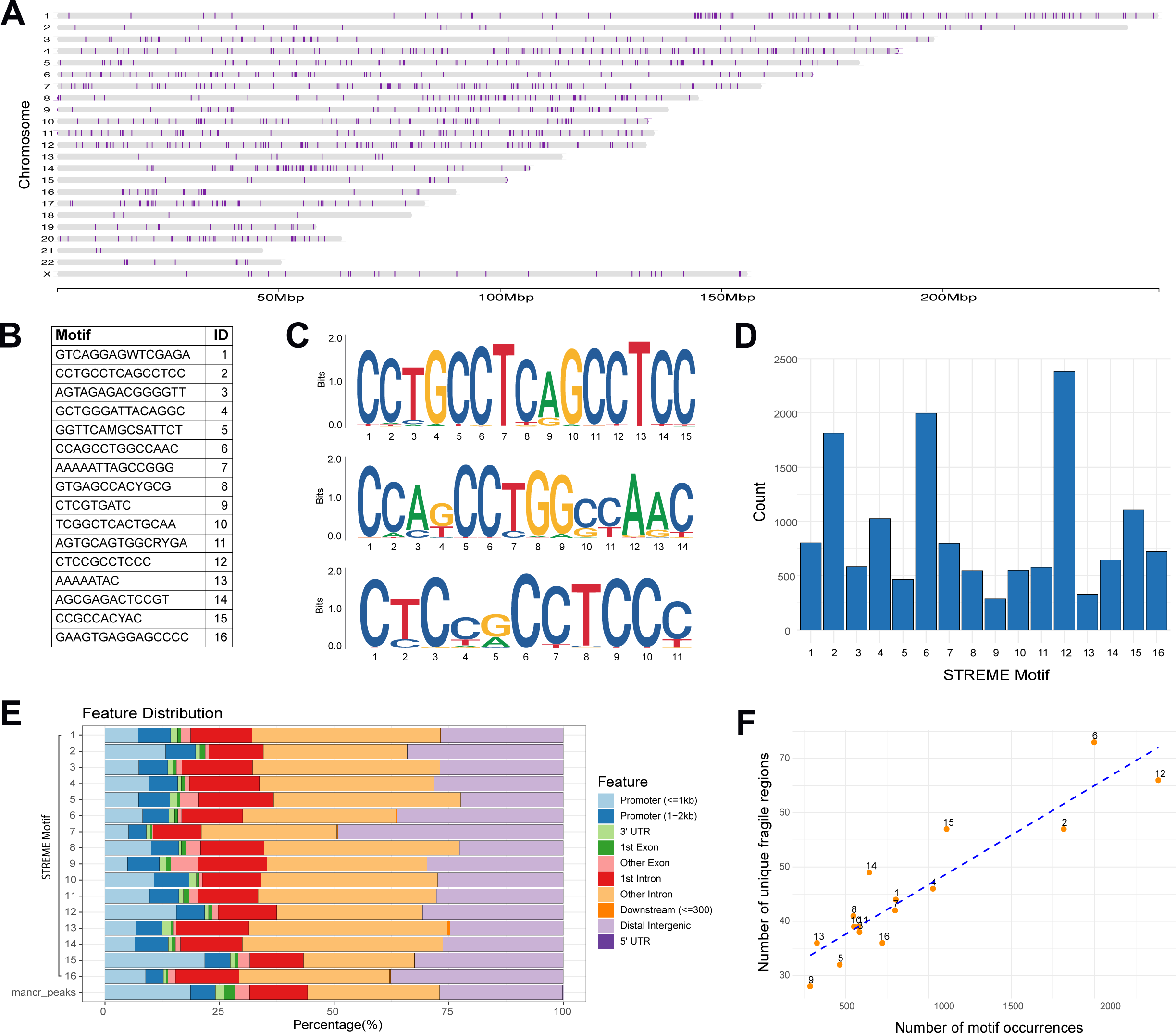
Identification of DNA MANCR-binding motifs. (A) MANCR-binding hotspots identified by ChIRP. Star represents location of MANCR on chromosome 10. (B) Sequences of 16 MANCR-binding DNA motifs and motif ID (C) Logo of top three motifs predicted by STREME analysis of MANCR ChIRP. In order, motifs are 2, 6, and 12. (D) Total number of each predicted motif occurrence within MANCR ChIRP regions. (E) Gene annotation of each motif within ChIRP regions and annotation of total ChIRP peaks. (F) Motif enrichment in fragile regions.

In addition to P53 binding sites, we identified genome-wide MANCR binding regions across each chromosome and identified MANCR interaction sites on all chromosomes (Fig 5A). The MANCR gene locus resides on chromosome 10, and there are a substantial number of MANCR interacting sites on this chromosome. However other chromosomes [19, 16, 22, and 13] have a greater number of MANCR binding sites (Fig 5A). DNA sequences associated with chromosomal MANCR-binding regions were analyzed by Simple, Thorough, Rapid, Enriched Motif Elicitation (STREME) (44) and identified 16 unique MANCR-binding DNA motifs (p-value < 0.05), which have an average GC content of 59.22% (Fig 5B). Identified motifs were localized to the center of the ChIRP peaks, and each motif occurred only once in the majority of MANCR-associated regions (Supp. Fig. 4). We found that 94% of the 1250 ChIRP peaks contained at least one predicted motif, and 86% of ChIRP peaks contained at least two of the identified motifs. This would strongly suggest there is a definite consensus component to these specific motifs that are associated with MANCR interaction. Each of the motifs is represented in at least 24% of ChIRP regions. Motif 12 is a GC-rich (81.82% GC) sequence that was found in the majority (68.48%) of the ChIRP peaks, followed by motifs 6 (55.92% of ChIRP peaks, 71.53% GC) and 2 (61.12% of ChIRP peaks, 73.33% GC) (Fig 5C-D). These data suggest that MANCR genomic binding sites are predominantly GC rich and that MANCR interaction with genomic DNA involves GC interactions.

Genomic regions associated with ChIRP-defined sites were analyzed using MANCR-associated motifs and defined subgenomic annotation. Identified MANCR-associated motifs within ChIRP regions were characterized and displayed comparable genic distributions (Fig 5E). They were enriched in introns and distal intergenic regions. Approximately 20% of MANCR binding sites were identified in promoter regions, suggesting that MANCR has an active role in gene regulation at specific promoters. A large percentage (30.17%) of binding sites are identified in intergenic regions. A total of 16% of MANCR binding sites are within defined fragile regions (45). A fragile region on chromosome 7, FRA7J, contained the highest number (313) of MANCR motif regions (Supp. Fig. 5A). Although the number of total motif-fragile region overlaps increases linearly by overall number of motifs, the intersection between motifs and unique fragile regions demonstrates unique enrichment of motifs 6 across fragile regions (Fig 5F). Motif 12 is associated with RNA Polymerase II promoters (BP), and transcription factor activity (MF) (46). Motifs 6 and 2 are both associated with translational elongation (BP) (46).

Having identified MANCR-binding sequences by ChIRP analysis, we determined the specific MANCR isoforms that preferentially bind to one or all of these motifs. Analysis of individual MANCR isoforms by homology prediction identified multiple complementary sequences of defined de novo motifs within MANCR sequences. The MANCR-201, MANCR-202, MANCR-203 and MANCR-205 isoforms are identified to bind the de novo MANCR-associated DNA motifs (p-value < 0.05), with MANCR-205 exhibiting a higher enrichment (e-value 7.1e-30) and increased potential binding sites (17 sites) compared to the other isoforms (Fig 6A). In contrast, significant DNA motif binding was not associated with MANCR-204, - 206, and -207. The complementary DNA-binding regions of MANCR-201 and -205 are located within the lncRNAs in both structured and unstructured regions (Fig 4B-C, Supp. Fig. 5B). Motifs 2, 6 and 12 are highlighted in red and are present primarily in regions predicted to form helices (Fig 4B-C). The distal region of MANCR-205 is densely enriched in DNA motif-binding regions and contains binding domains for 11 motifs, including motifs 2, 6, and 12 (Supp. Fig. 5B). Isoforms 201, 202, and 203 contain isolated domains of multi-motif-binding regions.

**Fig. 6.**
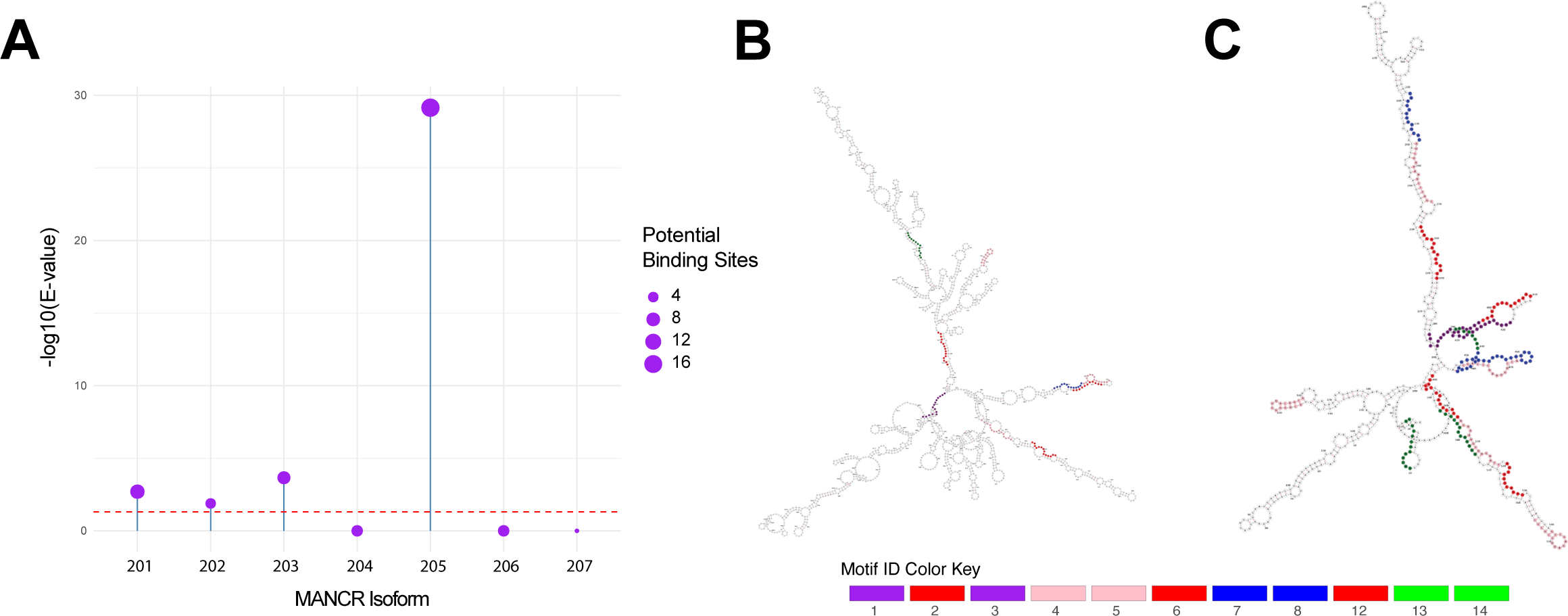
MANCR isoforms bind DNA with varying affinity. (A) Lollipop plot of e-values and number of predicted DNA-binding sites within each MANCR isoform. (B) Structural representation of DNA-binding regions within MANCR-201. (C) Structural visualization of predicted DNA-binding regions of MANCR-205. Motifs represented by colors by motif ID (Fig 3B).

### MANCR isoforms interact with specific miRNAs

A potential function for MANCR is interaction and sequestration of miRNAs. RIsearch2 identified 270 miRNAs with potential to interact with MANCR (Fig 7A). MANCR isoforms - 201, -204, and -206 were predicted to exhibit recognition and binding to similar miRNAs, with shared miRNA binding domains within MANCR (Fig 7B, D). The regions of MANCR-201, -204, and -206 identified as sites of miRNA binding indicate that some miRNA binding sites overlap with DNA binding sites (Fig 7D, Supp. Fig. 5B). However, MANCR-201 displays extensive miRNA binding towards the distal end of Arm 3 (around base 1, 000), which is not predicted to bind DNA, suggesting that a majority of miRNA binding occurs independently from DNA interactions (Fig 7D, Supp. Fig. 6A, Fig 2A-B, Fig 6B, Supp. Fig. 5B). MANCR-202, -203, - 205, and -207 isoforms appeared to predominantly bind unique miRNAs, with overlap between MANCR-202 and -203, with limited similarities in miRNA binding domains across isoforms (Fig 7B, D). The top ten miRNAs, which exhibited the highest number of unique interactions with MANCR across isoforms, each bound to at least four MANCR isoforms (Fig 7C, Supp. Fig. 6B). miRNA 6756-5p was identified as the miRNA with the highest number of unique MANCR-binding sites, and exhibited interactions with MANCR-201 in four unique locations across the lncRNA in both structured and unstructured regions with limited overlap with one DNA-binding domain (Fig 7C, E). This miRNA is known to interact with the Androgen Receptor (AR). Biological relevance is supported by decreased AR expression (RNA-seq) following MANCR knockdown. Regulatory consequences of predicted MANCR sponging are reinforced by decreased expression of over 70% of differentially expressed gene targets of the top ten identified miRNAs (p-value < 0.05) (Fig 7F).

**Fig. 7.**
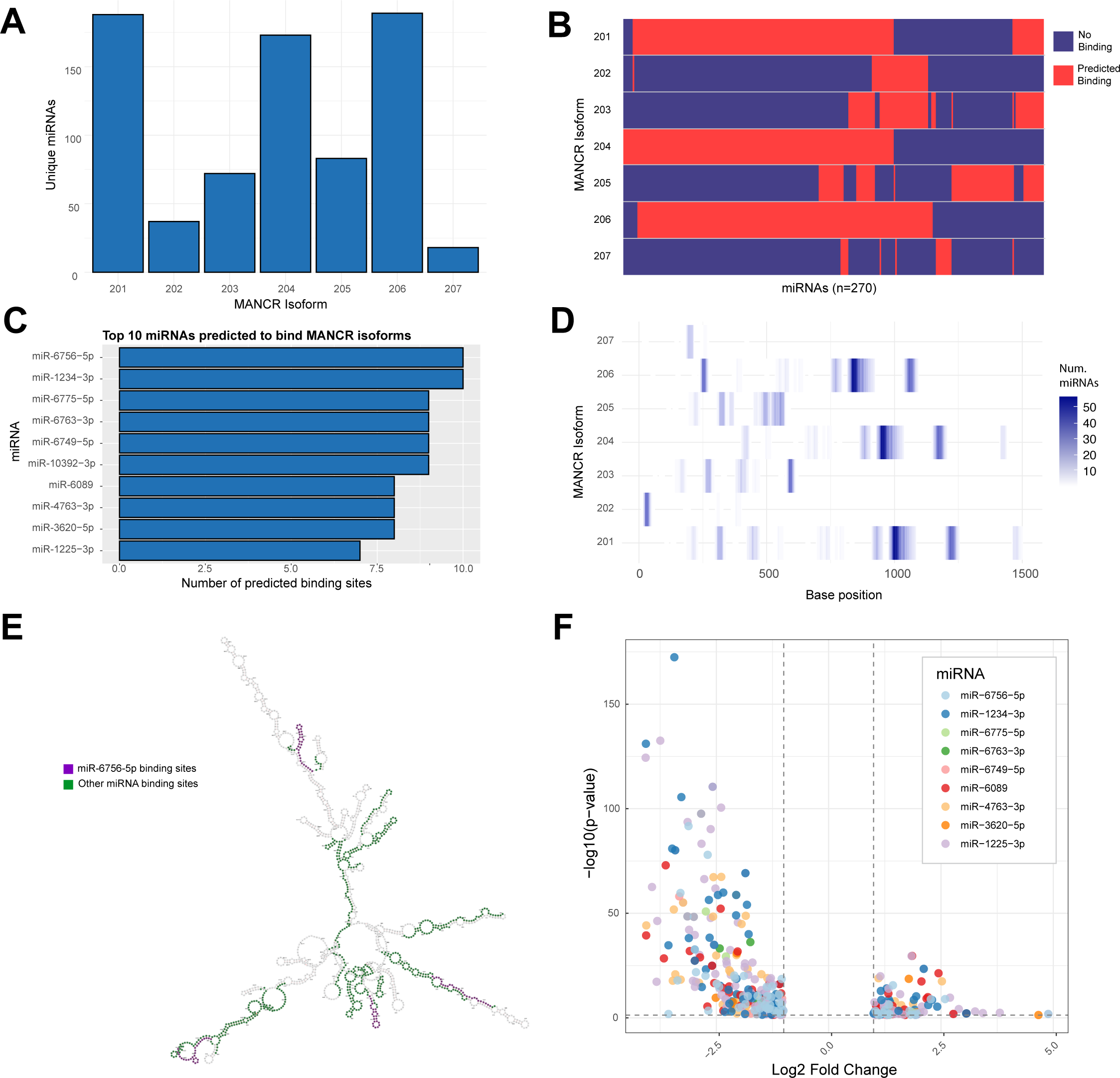
MANCR may sponge miRNAs associated with tumorigenesis and cancer progression. (A) Number of unique miRNAs binding to each MANCR isoform. (B) Clustered plot of miRNAs binding MANCR isoforms. Each red bar represents miRNA binding to each MANCR isoforms. Blue bars indicate the miRNA is not predicted to bind to the MANCR isoform. (C) Top ten miRNAs with the highest number of unique binding sites on all MANCR isoforms. (D) Heatmap of miRNA-binding hotspots on each MANCR isoforms. (E) Structural representation of miRNA binding regions of MANCR-201. Sites of predicted binding for hsa-miR-6756-5p are highlighted in purple, sites of all other predicted miR binding are highlighted in green. (F) Volcano plot of differential expression following MANCR knockdown of miRNA target genes.

## Discussion

In this study we identified computationally derived structures for the long non-coding RNA MANCR that provide insight into putative engagement of this oncolinc in mechanisms that mediate cancer-compromised biological control in aggressive breast cancer cells. We identified that seven MANCR isoforms have varied abundance, and this abundance changes based on cellular context that include hypoxia, drug resistance, and EMT. These isoforms have structural variation that confer several specific functions. We identified two consensus structures that are exhibited by the majority of isoforms. The propensity of the MANCR isoforms binding to protein was exhibited by computationally predicted and experimentally confirmed interactions with P53. Genomic interactions by MANCR were established, which include binding to intragenic binding sites as well as promoters that regulate P53 target-gene activity and miRNAs. MANCR regulation of genome stability is reinforced by direct evidence presented for motif-specific binding at fragile chromosomal sites and interactions with P53 and microRNAs.

Based on our findings, we identified that different MANCR isoforms engage in diverse roles during cancer progression. MANCR-205 expression in breast cancer is increased under both hypoxic conditions and following loss of the epithelial/tumor suppressor transcription factor RUNX1, suggesting MANCR isoform expression changes in response to cellular context or condition. This suggests that the switch in MANCR isoform expression changes the primary function of MANCR within the cell. This is important as MANCR-205 is enriched in DNA-binding motifs that enable it to bind genomic DNA. Upregulation of this isoform may lead to increased DNA binding and stabilization of the genome to allow for cancer progression. MANCR-205 upregulation following treatment of TNBC cells with NAMI-A, an antimetastatic drug that binds collagen, supports that this specific isoform of MANCR is upregulated in relation to cell survival and ECM remodeling (47). Our computational modeling of individual MANCR isoforms demonstrates unique biologically relevant functionality of isoform structural groups and domains that may bind proteins and nucleic acids independently.

This analysis furthers our understanding of how the structure of lncRNAs contributes critically to their functional characterization. Studies examining Metastasis-Associated Lung Adenocarcinoma Transcript 1 (MALAT1), a nuclear-localized lncRNA with complex oncogenic roles structural studies identified pseudoknots, structured tetraloops, and structured internal loops as well as intramolecular long-range interactions (11). In our analysis of MANCR, we identified 3D packing of hairpin domains, internal loops and stem structures. However, unlike MALAT1 our analysis did not identify pseudoknots or tetraloops. As observed for MALAT1, we found some intramolecular long-range interactions that are largely conserved across isoforms. MALAT1 is frequently mutated in cancer and these mutations are predicted to alter the structure of MALAT1 thereby exposing binding sites for microRNAs (48). Although no cancer-specific mutations in MANCR have yet been defined, it is clear that isoform-specific interactions may play a role in cancer progression based on transcript abundance and structural variation of isoforms. Recently, MALAT1 has been shown to bind to RNPS1, colocalize to nuclear speckles, and regulate alternative splicing (18). Similarly, MANCR has been shown to interact with hnRNP, an RNA splicing factor sometimes associated with nuclear speckles (14). In addition, we have identified that MANCR is associated with chromatin during mitosis (13). This interaction suggests that while both MANCR and MALAT1 have a role in oncogenesis, they are playing distinct and novel functional roles. Studies have identified stable loop structures in MALAT1 through RNPS1 binding, which allowed therapeutic targeting and destruction of this complex (18). This demonstrates the potential for specific cancer therapies targeting MANCR and highlights the importance of computational strategies to identify these targets. Our comprehensive examination of MANCR structure and molecular interactions provides understanding for potential molecular mechanisms of this cancer-associated lncRNA and indicates novel therapeutic strategies to disrupt these molecular interactions.

We predicted secondary and tertiary structures of seven MANCR isoforms with unique DNA, miRNA, and P53 protein binding interactions. We identified that MANCR isoforms vary widely in predicted structure, but contain conserved conformations in their trunk domains. MANCR-201 and MANCR-205 are the predominant isoforms in the majority of cancer cell lines, with diverse functions in DNA, protein and miRNA binding. MANCR-201, -204 and -206, share structural similarity and are predicted to sponge miRNAs in a similar manner. However, the predicted DNA binding functions of MANCR-201 appear to be dissimilar to MANCR-204 and -206. MANCR-205 is another predominant isoform of MANCR in cancer cell lines and shares modest structural similarity with MANCR-202 and -203. These three isoforms are predicted to bind DNA, with MANCR-205 containing the highest number of predicted DNA binding motif regions. The unique distribution of MANCR isoforms in each tissue and cell type may contribute to unique impacts of MANCR. Also, MANCR isoform expression is affected by external factors such as hypoxia, anticancer therapies, and oncogenic drivers. MANCR-205 is the most responsive to cellular changes and has the ability to interact with DNA, RNA, and P53, suggesting a diverse functionality in response to the cancer cellular environment. This would suggest that the diversity of functionality in MANCR isoforms may be a unique mechanism by which MANCR mediates pro-oncogenic effects.

Cancers with high MANCR expression and MANCR amplification, including lung cancer, head and neck cancer, ovarian cancer, and breast cancer also have high percentages of TP53 mutations (30). In HNSCC, mutated TP53 was associated with increased P53 expression, altered DNA binding affinity, altered interactions with substrates, and subsequently poor patient prognosis (49). These genetic lesions have been proposed as driver mutations in HNSCC and potential novel biomarkers for targeted therapeutic strategies (49). Across multiple cancers, MANCR amplification and/or expression is significantly correlated with P53 mutation. In breast cancer patients, high P53 expression is associated with P53 mutations in 70% of patients. This would strongly suggest a relationship of altered P53 function with MANCR expression and may indicate disruption of physical interaction. The arm 2 domain of MANCR-205 is predicted to interact with P53 tetramers both with and without the presence of DNA. Through RNA immunoprecipitation we demonstrate interactions between MANCR and P53, particularly with MANCR-205. Knockdown of MANCR resulted in decreased expression of P53-target genes and P53-mediated luciferase transcriptional activity. Although there are numerous P53 mutations in patients with high MANCR expression, the MDA-MB-231 cell line used for our analysis has a R280K mutation in the P53 protein. However, this did not prohibit MANCR interactions in gene expression analysis or luciferase activity. These data, combined with our prediction that MANCR-205 strongly binds DNA, suggests that MANCR may stabilize the interactions between P53 and DNA, including for certain P53 mutations, to allow for sustained functionality of P53. Our analysis predicted the interacting domains between MANCR-205 and P53, which was supported by altered binding efficiency in the presence of a P53 mutation in the predicted binding domain. This experimentally supports the identification of the binding domains and reveals the potential to therapeutically interrupt MANCR-P53 interactions. There is a requirement to mechanistically characterize lncRNAs as a basis for utilization in targeted cancer therapies (50).

There are diverse and central roles for lncRNAs in tumorigenesis, cancer progression, and metastasis (50). Abnormal expression of lncRNAs is associated with cancer cell proliferation and invasion, and lncRNAs have been identified as potential biomarkers for diagnosis, prognosis, and personalized treatments (50). The roles of lncRNAs in cancer are complex, and lncRNAs have both oncogenic and tumor-suppressive functions (3). Oncogenic lncRNAs have been associated with EMT, cancer stem cell formation, and angiogenesis. Identification of lncRNAs has the potential to support early cancer detection and improved prognosis for metastasis and recurrence (3). In breast cancer, lncRNA BCRT1 (breast cancer-related transcript 1) is elevated, and expression levels were reported to rise under hypoxic conditions, as was observed for MANCR (50). Knockdown of BCRT1 reduced metastasis and tumor expansion both in vitro and in vivo. A mechanistic explanation for BCRT1 activity may be the presence of Hypoxia-Inducible Factor-1α (HIF-1α) and Hypoxia-Responsive Elements (HREs) in the promoter region of BCRT1, linking this lncRNA to a pivotal pathway in breast cancer progression (50). Increased expression of MANCR under hypoxic conditions across cancer cell lines provokes further investigation into the relationship between MANCR and HREs. Tang et al suggest MANCR as a novel therapeutic target for HNSCC due to the correlation between MANCR expression and immune response (51).

Transcriptome analysis revealed MANCR-depletion was associated with pre-mRNA splicing, controlling DNA repair, and differential expression of genes involved in cell cycle regulation, ECM-receptor interaction, and checkpoint response (51–53). Through positive regulation of NET1A, which regulates metastatic processes and DNA damage checkpoint control, MANCR has been implicated in modulating transcription factor binding and promoting metastasis (52). MANCR knockdown in HNSCC resulted in 224 differentially expressed genes demonstrating that MANCR plays a role in gene expression in these cells (51). Our previous findings suggest that MANCR regulates over 2, 000 genes in MDA-MB-231 TNBS cells (13). Predicted MANCR DNA motifs and experimentally observed MANCR DNA binding regions are enriched at introns and distal intergenic regions of genes associated with transcription and translation. MANCR motifs and experimentally-derived regions identified by ChIRP are enriched in established genome fragile regions, suggesting a potential role in genome stability in both genic and distal intergenic regions, in addition to potential associations with gene regulation. Singh et al focus on MANCR-201, which interacts with motifs 1, 4, and 6 that are enriched in translation and RNA processing (14). MANCR-205 also binds these motifs, as well as eight additional motifs. Much of the apex, trunk, and arm 1 of MANCR-205 are predicted to bind DNA. Consideration of MANCR-205 will be important for experiments on MANCR-mediated control of DNA stability and repair as well as transcription.

Another level of MANCR-mediated transcriptional regulation is sponging of miRNAs by MANCR. In this study we provide evidence that MANCR isoforms bind to a cohort of miRNAs. The top predicted miRNA is miR-6756-5p, which regulates Androgen Receptor (AR). Following loss of MANCR, AR expression decreases in MDA-MB-231 cells, suggesting a functional link between MANCR and AR expression mediated by miR-6756-5p. In parallel, we demonstrate that MANCR physically interacts with P53 justifying inclusion within the P53 regulatory network. Previous studies have shown that breast tumors with mutant P53 display decreased AR expression and increased EMT with poorer clinical outcomes (22). These observations are consistent with MANCR control of AR expression through transcriptional regulation with P53 and miRNA-mediated post-transcriptional mechanisms. Further investigation into MANCR competency to fine-tune gene expression could provide insights into DNA-damage response, EMT, and cancer progression.

A recent study of TNBC utilizing MDA-MB-231 and Hs578t cell lines determined a mechanism for Androgen Receptor (AR) functionality in promoting tumorigenesis in TNBC was through transcriptionally activating long noncoding RNA SOX2-OT (54). Subsequently, SOX2-OT sponges miR-320a-5p, which is a regulator of CCR5. Molecular sponging of miR-320a-5p by SOX2-OT has been shown to contribute to increased cell proliferation and inhibition of apoptosis in TNBC in vitro, and TNBC tumorigenesis in vivo. An analysis of tissue microarray and data from The Cancer Genome Atlas supported this tumorigenesis mechanism in breast cancer patients (54). Our data similarly predict miRNA sponging as a mechanism for MANCR, and AR is a target of our top predicted miRNA, which is miR-6756-5p. Decreased expression of AR is observed through RNA-seq following MANCR knockdown, supporting a functional role for MANCR in regulating AR through miRNA sponging. Further, MANCR is predicted to interact with miRNAs in an isoform-specific manner. MANCR-201, -204, and -206 bind hundreds of miRNAs across structured and unstructured regions. These isoforms contain miRNA-binding hotspots that are distinct from their DNA-binding domains. The other MANCR isoforms bind largely unique miRNAs with limited overlaps in binding regions across isoforms. Upon MANCR degradation, we anticipate freeing of miRNAs and subsequent downregulation of other miRNA targets. As expected, the majority of miRNA-target genes that are differentially expressed following MANCR knockdown are significantly downregulated. Although direct miRNA sponging by MANCR has not been experimentally validated, these data support a model in which MANCR modulates gene expression post-transcriptionally through miRNA-dependent regulation to promote tumorigenesis.

These findings support a model in which MANCR operates as a functionally complex oncolnc lncRNA, and isoform diversity contributes to context-dependent regulatory interactions. Conserved structural features suggest core functional elements that interact with transcriptional regulatory factors and DNA while allowing structural and sequence variation that modulates binding specificity. The identification of MANCR interactions with P53, fragile genomic regions, and miRNAs further supports a role for this lncRNA in genomic stability and transcriptional regulation. These analyses integrate isoform variation, RNA structure, and molecular interactions to provide a structural and functional framework for understanding MANCR function in cancer cells. Further studies interrogating the MANCR-chromatin interactions and MANCR-mediated transcriptional regulation will be critical to elucidate the role of this lncRNA throughout cancer development and progression.

## Materials and Methods

### cBioPortal Investigation of Patient Survival and MANCR and TP53 Alterations

Patient survival data from Pan-cancer Analysis of Advanced and Metastatic Tumors (30) was surveyed for 86 samples with high MANCR expression (above 0.5) and 86 samples with low MANCR expression. A Kaplan-Meier plot was generated by CBioPortal (55) to visualize survival for each group and p-value and hazard ratio were calculated.

The association between TP53 mutation and MANCR (Mitotically-Associated Long Non-Coding RNA) amplification and expression across cancer types was analyzed through cBioPortal (55) for the PanCancer ICGC/TCGA Nature 2020 dataset (30). Cancer Type Summary plots were generated for TP53 and MANCR, and Mutual Exclusivity was calculated by Fisher’s Exact Test, p < 0.05, to determine co-occurrence of TP53 mutation and MANCR copy number amplification.

Within breast cancer, we evaluated data from 33 studies available on cBioPortal (55) with data on TP53 mutations and MANCR copy number variations and increased expression. Mutual Exclusivity was calculated by Fisher’s Exact Test to determine co-occurrence of TP53 mutation and MANCR amplification within these breast cancer studies. The Plots module on cBioPortal (55) was used to obtain a detailed overview of TP53 mutations within breast cancers with MANCR CNVs and high expression.

### Cell Culture and Treatment

MDA-MB-231 cells were grown as previously described (13). Briefly, MDA-MB-231 cells are obtained from ATCC and cultured in DMEM/F12 supplemented with 10% fetal bovine serum (FBS) and 1% penicillin/streptomycin and 1% L-Glutamine. A pool of two anti-sense Gapmers was designed to knock down all MANCR isoforms. MDA-MB-231 cells were treated with the Gapmer pool or with a negative control ASO for 24 hours in DMEM/F12 supplemented with 10% fetal bovine serum (FBS) and and 1% L-Glutamine for RNA-seq.

### RNA Sequencing Analysis of MDA-MB-231 Cells

RNA from three replicates of MDA-MB-231 cells was collected following 24 hours of gapmer treatment and sequenced at 30x depth. Fastq files were trimmed using Trim Galore v0.6.6 (56) and aligned to GRCh38.p14 by STAR v2.7.11b with default parameters (57). DESeq2 performed differential gene expression analysis with FC > |1| and FDR < 0.05 statistical cut-offs (58).

For pathway analysis, differentially expressed genes were compared to the P53 KEGG pathway in R (v4.3.2) (40). Downregulated TP53 targets were visualized and modeled using Cytoscape (v3.10.3) (59).

### Secondary and Tertiary Structure Prediction of MANCR

Predicted secondary structures of MANCR isoforms were generated using RNAalifold and Alidot (34), for both individual isoforms and multiple sequence alignments utilizing consensus sequences. Base pairing probabilities were computed with RNAfold -p for each MANCR isoform (34). Multiple sequence alignments (MSA) were generated by MAFFT (60) and subsequently submitted to RNAalifold with -p, -aln and -sci flags to compute base pairing probabilities, minimum free energy (MFE), and structure conservation index (SCI). Alidot was used to identify structurally conserved stems within MSAs (34). Custom scripts were written in Python (v 3.9.21) to calculate mean pairwise identity. Selected structural domains were modeled in 3D using RNAComposer (61).

### Luciferase Assay

Gapmers targeting MANCR, P53, and a negative control ASO were generated by Qiagen (Supp Table 3). MANCR required a pool of gapmers to target all seven isoforms. The same MANCR gapmers were used in RNA-seq. MDA-MB-231 cells were treated with corresponding gapmers and the PG13-luc luciferase plasmid from Addgene, which contains a P53 binding motif followed by the luciferase gene. PG13-luc (wt p53 binding sites) was a gift from Bert Vogelstein (Addgene plasmid # 16442 ; http://n2t.net/addgene:16442 ; RRID:Addgene_16442) (62). For three replicates, cells were incubated with PG13-luc and gapmers for 24 hours and then assayed following the Promega Dual-Luciferase Reporter Assay kit E1910 for three replicates. Luciferase luminescence was normalized by Renilla luminescence. Fold change relative to the negative control was calculated for the normalized mean values. Statistical significance was evaluated using Welch’s two-sample t-test between each experimental condition compared to the negative control. Data was graphed by ggplot2 (63) as mean value with error bars of SEM, with significance defined as p < 0.05.

### Chromatin Isolation by RNA Purification (ChIRP)

MANCR interactions with genomic sequences were established by ChIRP in MDA-MB-231 cells. Biotinylated antisense oligonucleotides for MANCR isoforms 201-207 were designed with the Biosearch Technoligies ChIRP Probe Designer and purchased from there. For two replicates, cells were crosslinked with 1% glutaraldehyde followed by 3% formaldehyde and the crosslinking was quenched with glycine. The chromatin was isolated and sheared by sonication and all 22 MANCR probes were pooled and allowed to hybridize with the MANCR-Chromatin regions overnight. NGS libraries were prepared from the hybridized regions of chromatin that were isolated by the MANCR probes and sequenced at 75G per sample at Novogene. Enriched genomic DNA was purified and sequenced by Novogene. Fastq files were trimmed using Trim Galore v0.6.6 and aligned to GRCh38.p14 by STAR v2.7.11b with default parameters (56, 57). Peaks were called by macs3 (64) and ENCODE black-list regions (65) were removed to identify MANCR-associated regions. HOMER (41)was used to call motifs within MANCR-associated regions and identify potential proteins interacting with MANCR on chromatin. FIMO (42) was used to confirm enrichment of the P53 motif within MANCR-associated regions.

### Motif Discovery and Functional Annotation

To identify MANCR motifs, we used STREME (part of the MEME Suite Tools) for de novo motif discovery from MANCR-ChIRP data (44). Motif annotation was performed with GOMo (46), linking identified motifs to biological processes. Genomic binding positions were annotated using TxDb (v3.7.0) for hg38 in R. MAST (46) was employed to evaluate enrichment of the identified motifs within MANCR isoforms.

### Protein Interaction Modeling

The AlphFold Server (35) was utilized to screen interactions between P53 tetramers and each structural domain of MANCR-205. Arm 2 of MANCR was identified as the predicted interacting domain between MANCR and P53 tetramers. Due to the unique needs of RNA folding prediction compared to protein predictions, MANCR:P53 interaction was confirmed by HDOCK (36). The 3D structure of MANCR was predicted using RNAComposer (61) for structural domains of MANCR isoform 205. The RNAComposer 3D structures and publicly available P53 structures (PDBIDs: 2OCJ and 2ADY) were modeled using HDOCK, a hybrid docking algorithm combining template-based modeling with free docking (36). The predicted structures were visualized in PyMOL and residues closer than 3 Å were identified to characterize direct interaction.

### RNA Immunoprecipitation (RIP) and qPCR

RIP-qPCR was conducted with three replicates to validate MANCR interaction with P53 protein. P53-RIP complexes were obtained in MDA-MB-231 cell lysates through immunoprecipitation using anti-p53 antibody (clone pAb122, Abcam, ab90363) and IgG (FisherScientific, catalog no. NC0411739), and co-precipitated RNA was quantified by RT-qPCR using MANCR-specific primers. TBP and MALAT1 primers were used as positive controls, and SMARCE1 primers were used as a negative control. SMARCE1 primers were previously validated. Values were normalized to input, and enrichment of RNAs in P53 pull-down was calculated over IgG. Statistical significance was assessed using ΔCt values relative to IgG controls with Benjamini–Hochberg correction.

### RIP-Seq

RNA from P53-RIP complexes described above were sequenced to profile P53-associated transcripts, with an IgG-RIP as a background control. Data from our RIP-seq experiments were complemented by publicly available RIP-seq datasets for p53 from NCBI Sequence Read Archive associated with BioProject PRJNA1210463. Reads were processed using the nextflow rnaseq pipeline (66), and P53-bound RNAs were identified as RNAs with at least one isoform with Log2FC > 0.5, p-value < 0.05, and TPM > 1.

### miRNA Binding Prediction

The mature.fa file of mature miRNA sequences was downloaded from miRBase for analysis (67). To identify miRNAs that may bind MANCR, we used RIsearch2 (68) with an energy cutoff of -25 kcal/mol. Candidate miRNAs were ranked in R based on the number of unique predicted binding sites along all MANCR isoforms and visualized using ggplot2 (63). Gene targets of miRNAs were identified by multiMiR and differential expression of the miRNA gene targets were identified in R from RNA-seq with and without MANCR gapmer knockdown (69).

## Supplementary Materials

**Supp. Fig. 1**. Hypoxic conditions alter MANCR expression. (A) MANCR isoform expression in MDA-MB-436 expression under normal and hypoxic conditions. (B) MANCR isoform expression in MDA-MB-157 expression under normal and hypoxic conditions. (C) MANCR isoform expression in HCC70 expression under normal and hypoxic conditions.

**Supp. Fig. 2.** Predicted MFE conformations for MANCR isoforms MANCR-202 (A), -203 (B), - 204 (C), -206 (D), and -207 (E). Color scale represents base pairing probability for minimum free energy (MFE) structure with red indicating high base pairing probability and blue indicating low base pairing probability.

**Supp. Fig. 3**. MANCR binds P53. (A) Predicted interaction between the P53 structure (PDBID: 2OCJ, not including DNA) and the predicted structure of MANCR-205 Arm 2. (B) Average TPM of MANCR isoforms pulled down by WT-P53 and P53R175 mutant RIP-seq.

**Supp. Fig. 4**. DNA-binding MANCR motifs predicted by STREME based on MANCR ChIRP-seq.

**Supp. Fig. 5**. DNA-MANCR interactions are isoform specific. (A) Fragile sites containing MANCR DNA-binding sites motifs show enrichment in FRA7J. (B) DNA-binding regions of MANCR along the length of each isoform predicted by MAST (46).

**Supp. Fig. 6**. Binding of miRNAs to MANCR isoforms. (A) Number of miRNAs that bind across the MANCR-201 isoform depicted as a line graph. (B) Number of MANCR isoforms bound by the topo ten identified miRNAs.

## REFERENCES

1. De Martino, M., Esposito, F., and Pallante, P. (2021) Long non-coding RNAs regulating multiple proliferative pathways in cancer cell Transl Cancer Res 10, 3140–3157

2. Alzarea, S. I. (2025) Non-coding RNA-mediated gene regulation in Alzheimer’s disease pathogenesis: molecular insights and emerging innovations Saudi Pharm J 33, 33

3. Chainsee Saini, P. V., Simran Maharshi, Bhavika Baweja, Rajeev Nema (2025) Involvement of lncRNA in cancer diagnosis and prognosis and clinical implications Rep Pract Oncol Radiother

4. Zeng, Z., Zhou, F., Zheng, Y., Shi, W., Hu, W., Wang, X., et al. (2025) Comprehensive analysis of LncRNA-miRNA-mRNA CeRNA network associated with umbilical cord blood PBMC in down syndrome Scientific Reports 15,

5. Braga, E. A., Fridman, M. V., Burdennyy, A. M., Loginov, V. I., Dmitriev, A. A., Pronina, I. V. et al. (2023) Various LncRNA Mechanisms in Gene Regulation Involving miRNAs or RNA-Binding Proteins in Non-Small-Cell Lung Cancer: Main Signaling Pathways and Networks International Journal of Molecular Sciences 24, 13617

6. Pisignano, G., and Ladomery, M. (2021) Epigenetic Regulation of Alternative Splicing: How LncRNAs Tailor the Message Non-Coding RNA 7, 21

7. Song, Y. J., Shinn, M. K., Bangru, S., Wang, Y., Sun, Q., Hao, Q. et al. (2026) LncRNA-splicing factor condensates regulate hypoxia-responsive pre-mRNA processing near nuclear speckles Mol Cell 86, 1061–1080 e1010

8. Dalal, M., Joshi, R., Ajithkumar, P., Singh, J., Suri, V., Chatterjee, A. et al. (2025) Novel insights into hypoxia-driven transcriptomic and epigenetic landscapes in grade 3 meningioma Journal of Translational Medicine 24,

9. McCown, P. J. (2019) Secondary Structural Model of Human MALAT1 Reveals Multiple Structure–Function Relationships International Journal of Molecular Sciences

10. Mishra, A. (2024) Metastasis-Associated Lung Adenocarcinoma Transcript 1 (MALAT1) lncRNA Conformational Dynamics in Complex with RNA-Binding Protein with Serine-Rich Domain 1 (RNPS1) in the Pan-cancer Splicing and Gene Expression ACS Omega

11. Phillip J. McCown, M. C. W., Luc Jaeger 2ORCID and Jessica A. Brown 1 (2019) Secondary Structural Model of Human MALAT1 Reveals Multiple Structure–Function Relationships Int. J. Mol. Sci

12. Ma, L., Liu, X., Roopashree, R., Kazmi, S. W., Jasim, S. A., Phaninder Vinay, K., et al. (2025) Long non-coding RNAs (lncRNAs) in cancer development: new insight from STAT3 signaling pathway to immune evasion Clin Exp Med 25, 53

13. Tracy, K. M., Tye, C. E., Ghule, P. N., Malaby, H. L., Stumpff, J., Stein, J. L., et al. (2018) Mitotically-associated lncRNA (MANCR) Affects Genomic Stability and Cell Division in Aggressive Breast Cancer Molecular Cancer Research

14. Singh, D. K., Cong, Z., Song, Y. J., Liu, M., Chaudhary, R., Liu, D. et al. (2024) MANCR lncRNA Modulates Cell-Cycle Progression and Metastasis by Cis-Regulation of Nuclear Rho-GEF Mol Cell Biol 44, 372–390

15. Bone, M., and Inman, G. J. (2025) Alternative transcription increases isoform complexity in Long Non-Coding RNAs and alters their functions in cancer Noncoding RNA Res 14, 38–50

16. Abulwerdi, F. A., Xu, W., Ageeli, A. A., Yonkunas, M. J., Arun, G., Nam, H. et al. (2019) Selective Small-Molecule Targeting of a Triple Helix Encoded by the Long Noncoding RNA, MALAT1 ACS Chem Biol 14, 223–235

17. Grice, F. A. (2019) Selective Small-Molecule Targeting of a Triple Helix Encoded by the Long Noncoding RNA, MALAT1 ACS Chemical Biology

18. Aanchal Mishra, S. M. (2024) Metastasis-Associated Lung Adenocarcinoma Transcript 1 (MALAT1) lncRNA Conformational Dynamics in Complex with RNA-Binding Protein with Serine-Rich Domain 1 (RNPS1) in the Pan-cancer Splicing and Gene Expression ACS Omega

19. Liu, J., Zhang, C., Ma, B., Qi, J., Zhang, X., and Zhao, H. (2025) A ferroptosis-related risk model for non-coding RNA AC002331.1 in colon cancer: construction via competing endogenous RNA network analysis Discover Oncology 16,

20. Liu, Z. (2023) Long non-coding RNA DINO promotes cisplatin sensitivity in lung adenocarcinoma via the p53-Bax axis Journal of Thoracic Disease

21. Schmitt, C. (2021) p53-intact cancers escape tumor suppression through loss of long noncoding RNA Dino Cell Reports

22. Singh, A., Gupta, R., and Kulshreshtha, R. (2026) Capturing the Variations in Mutant p53-Driven Regulatory Networks in Breast Cancer Subtypes, Its Clinical Relevance and a Novel Association With Androgen Receptor and EMT IUBMB Life 78,

23. Song, J., Cui, Q., and Gao, J. (2024) Roles of lncRNAs related to the p53 network in breast cancer progression Frontiers in Oncology 14,

24. Sanchez-Marin, D., Trujano-Camacho, S., Perez-Plasencia, C., De Leon, D. C., and Campos-Parra, A. D. (2022) LncRNAs driving feedback loops to boost drug resistance: sinuous pathways in cancer Cancer Lett 543, 215763

25. Chaleshi, V., Irani, S., Alebouyeh, M., Mirfakhraie, R., and Asadzadeh Aghdaei, H. (2020) Association of lncRNA11p53 regulatory network (lincRNA11p21, lincRNA11ROR and MALAT1) and p53 with the clinicopathological features of colorectal primary lesions and tumors Oncology Letters 10.3892/ol.2020.11518

26. Jeffers, L. K., Duan, K., Ellies, L. G., Seaman, W. T., Burger-Calderon, R. A., Diatchenko, L. B. et al. (2013) Correlation of Transcription of MALAT-1, a Novel Noncoding RNA, with Deregulated Expression of Tumor Suppressor p53 in Small DNA Tumor Virus Models Journal of Cancer Therapy 04, 774–786

27. Kotake, Y., Kitagawa, K., Ohhata, T., Sakai, S., Uchida, C., Niida, H. et al. (2016) Long Non-coding RNA, PANDA, Contributes to the Stabilization of p53 Tumor Suppressor Protein Anticancer Res 36, 1605–1611

28. Liu, J., Ben, Q., Lu, E., He, X., Yang, X., Ma, J., et al. (2018) Long noncoding RNA PANDAR blocks CDKN1A gene transcription by competitive interaction with p53 protein in gastric cancer Cell Death & Disease 9,

29. Jing, Q., Chen, Q., Wang, G., Wu, T., Wang, L., Xiong, Q., et al. (2025) LINC02593 impedes cell senescence via COP1-mediated p53 degradation in cervical cancer Cell Signal 134, 111907

30. Pleasance, E., Titmuss, E., Williamson, L., Kwan, H., Culibrk, L., Zhao, E. Y. et al. (2020) Pan-cancer analysis of advanced patient tumors reveals interactions between therapy and genomic landscapes Nat Cancer 1, 452–468

31. Frankish, A., Carbonell-Sala, S., Diekhans, M., Jungreis, I., Loveland, J. E., Mudge, J. M. et al. (2023) GENCODE: reference annotation for the human and mouse genomes in 2023 Nucleic Acids Res 51, D942–D949

32. Fritz, A. J., Greenyer, H., Dillac, L., Chavarkar, P., Ullah, R., Malik, M., et al. (2026) Acute degron-mediated RUNX1 loss reprograms enhancer activity to epigenetically drive epithelial destabilization and initiate cancer hallmarks openRxiv,

33. Harris, A. R., Panigrahi, G., Liu, H., Koparde, V. N., Bailey-Whyte, M., Dorsey, T. H. et al. (2023) Chromatin Accessibility Landscape of Human Triple-negative Breast Cancer Cell Lines Reveals Variation by Patient Donor Ancestry Cancer Res Commun 3, 2014–2029

34. Lorenz, R., Bernhart, S. H., Höner Zu Siederdissen, C., Tafer, H., Flamm, C., Stadler, P. F., et al. (2011) ViennaRNA Package 2.0 Algorithms for Molecular Biology 6, 26

35. Abramson, J., Adler, J., Dunger, J., Evans, R., Green, T., Pritzel, A. et al. (2024) Accurate structure prediction of biomolecular interactions with AlphaFold 3 Nature 630, 493–500

36. Yan, Y., Tao, H., He, J., and Huang, S.-Y. (2020) The HDOCK server for integrated protein–protein docking Nature Protocols 15, 1829–1852

37. Farmer, G., Colgan, J., Nakatani, Y., Manley, J. L., and Prives, C. (1996) Functional Interaction between p53, the TATA-Binding Protein (TBP), and TBP-Associated Factors In Vivo Molecular and Cellular Biology 16, 4295–4304

38. Ibrahim, B., Gobran, M., Metwalli, A.-E., Abd Elhady, W., Tolba, A., and Omar, W. (2023) Interplay of LncRNA TUG1 and TGF-β/P53 Expression in Colorectal Cancer Asian Pacific Journal of Cancer Prevention 24, 3957–3968

39. Sun, G., Wang, Y., Zhang, J., Lin, N., and You, Y. (2018) MiR-15b/HOTAIR/p53 form a regulatory loop that affects the growth of glioma cells Journal of Cellular Biochemistry 119, 4540–4547

40. Kanehisa, M., Furumichi, M., Sato, Y., Ishiguro-Watanabe, M., and Tanabe, M. (2021) KEGG: integrating viruses and cellular organisms Nucleic Acids Res 49, D545–D551

41. Heinz, S., Benner, C., Spann, N., Bertolino, E., Lin, Y. C., Laslo, P. et al. (2010) Simple combinations of lineage-determining transcription factors prime cis-regulatory elements required for macrophage and B cell identities Mol Cell 38, 576–589

42. Grant, C. E., Bailey, T. L., and Noble, W. S. (2011) FIMO: scanning for occurrences of a given motif Bioinformatics 27, 1017–1018

43. Fischer, M., and Sammons, M. A. (2024) Determinants of p53 DNA binding, gene regulation, and cell fate decisions Cell Death & Differentiation 31, 836–843

44. Bailey, T. L. (2021) STREME: accurate and versatile sequence motif discovery Bioinformatics 37, 2834–2840

45. Kumar, R., Nagpal, G., Kumar, V., Usmani, S. S., Agrawal, P., and Raghava, G. P. S. (2019) HumCFS: a database of fragile sites in human chromosomes BMC Genomics 19,

46. Bailey, T. L., Boden, M., Buske, F. A., Frith, M., Grant, C. E., Clementi, L., et al. (2009) MEME SUITE: tools for motif discovery and searching Nucleic Acids Research 37, W202–W208

47. Bergamo, A., Gerdol, M., Lucafò, M., Pelillo, C., Battaglia, M., Pallavicini, A. et al. (2015) RNA-seq analysis of the whole transcriptome of MDA-MB-231 mammary carcinoma cells exposed to the antimetastatic drug NAMI-A Metallomics 7, 1439–1450

48. Yang, Q., Zheng, W., Shen, Z., Huang, G., and Yang, G. (2020) MicroRNA Binding Site Polymorphisms of the Long-Chain Noncoding RNA MALAT1 are Associated with Risk and Prognosis of Colorectal Cancer in Chinese Han Population Genet Test Mol Biomarkers 24, 239–248

49. Ashraf Attia Mahmoud, M. F. R., Edison Eukun Sage, Qurashi M. Ali, Omnia H. Suliman, Sabah A. E. Ibrahim, Osama Mohamed, Samar Abdelrazeg, Sofia B. Mohamed (2025) The impact of mutations on TP53 protein and MicroRNA expression in HNSCC: Novel insights for diagnostic and therapeutic strategies PLOS One

50. Zhicheng Li, D. W. X. Z. (2024) Unveiling the functions of five recently characterized lncRNAs in cancer progression Clinical and Translational Oncology

51. Jianfei Tang, M. B., Zhangui Tang (2022) Long-Noncoding RNA MANCR is Associated With Head and Neck Squamous Cell Carcinoma Malignant Development and Immune Infiltration Front. Genet.

52. Deepak K. Singh, Z. C., You Jin Song, Minxue Liu, Ritu Chaudhary, Dazhen Liu, Yu Wang, Rishabh Prasanth, Rajendra K C, Simon Lizarazo, Miriam Akhnoukh, Omid Gholamalamdari, Anurupa Moitra, Lisa M. Jenkins, Rohit Bhargava, Erik R. Nelson, Kevin Van Bortle, Supriya (2024) MANCR lncRNA Modulates Cell-Cycle Progression and Metastasis by Cis-Regulation of Nuclear Rho-GEF Molecular and Cellular Biology

53. Kirsten M Tracy 1, C. E. T., Prachi N Ghule 1, Heidi LH Malaby 2, Jason Stumpff 2, Janet L Stein 1, Gary S Stein 1, Jane B Lian 1 (2018) Mitotically-associated lncRNA (MANCR) Affects Genomic Stability and Cell Division in Aggressive Breast Cancer Mol Cancer Res

54. Hu, Y., Bian, J., Chen, W., Shi, J., Wei, X., Du, Y., et al. (2025) Androgen receptor-induced lncRNA SOX2-OT promotes triple-negative breast cancer tumorigenesis via targeting miR-320a-5p-CCR5 axis J Biol Chem 301, 108428

55. de Bruijn, I., Kundra, R., Mastrogiacomo, B., Tran, T. N., Sikina, L., Mazor, T. et al. (2023) Analysis and Visualization of Longitudinal Genomic and Clinical Data from the AACR Project GENIE Biopharma Collaborative in cBioPortal Cancer Res 83, 3861–3867

56. Andrews, S., Krueger, F., Segonds-Pichon, A., Biggins, L., Virk, B., Dalle-Pezze, P., et al. (2015) Trim galore Trim Galore

57. Dobin, A., Davis, C. A., Schlesinger, F., Drenkow, J., Zaleski, C., Jha, S. et al. (2013) STAR: ultrafast universal RNA-seq aligner Bioinformatics 29, 15–21

58. Love, M. I., Huber, W., and Anders, S. (2014) Moderated estimation of fold change and dispersion for RNA-seq data with DESeq2 Genome Biology 15,

59. Shannon, P., Markiel, A., Ozier, O., Baliga, N. S., Wang, J. T., Ramage, D. et al. (2003) Cytoscape: A Software Environment for Integrated Models of Biomolecular Interaction Networks Genome Research 13, 2498–2504

60. Katoh, K., Rozewicki, J., and Yamada, K. D. (2019) MAFFT online service: multiple sequence alignment, interactive sequence choice and visualization Briefings in Bioinformatics 20, 1160–1166

61. Sarzynska, J., Popenda, M., Antczak, M., and Szachniuk, M. (2023) RNA tertiary structure prediction using RNAComposer in CASP15 Proteins: Structure, Function, and Bioinformatics 91, 1790–1799

62. el-Deiry, W. S., Tokino, T., Velculescu, V. E., Levy, D. B., Parsons, R., Trent, J. M., et al. (1993) WAF1, a potential mediator of p53 tumor suppression Cell 75, 817–825

63. Wickham, H. (2016) ggplot2: Elegant Graphics for Data Analysis, Springer-Verlag New York,

64. Zhang, Y., Liu, T., Meyer, C. A., Eeckhoute, J., Johnson, D. S., Bernstein, B. E. et al. (2008) Model-based Analysis of ChIP-Seq (MACS) Genome Biology 9, R137

65. Amemiya, H. M., Kundaje, A., and Boyle, A. P. (2019) The ENCODE Blacklist: Identification of Problematic Regions of the Genome Sci Rep 9, 9354

66. Ewels, P. A., Peltzer, A., Fillinger, S., Patel, H., Alneberg, J., Wilm, A. et al. (2020) The nf-core framework for community-curated bioinformatics pipelines Nature Biotechnology 38, 276–278

67. Kozomara, A., Birgaoanu, M., and Griffiths-Jones, S. (2019) miRBase: from microRNA sequences to function Nucleic Acids Research 47, D155–D162

68. Alkan, F., Wenzel, A., Palasca, O., Kerpedjiev, P., Anders, Stadler, P. F., et al. (2017) RIsearch2: suffix array-based large-scale prediction of RNA–RNA interactions and siRNA off-targets Nucleic Acids Research 10.1093/nar/gkw1325gkw1325

69. Ru, Y., Kechris, K. J., Tabakoff, B., Hoffman, P., Radcliffe, R. A., Bowler, R. et al. (2014) The multiMiR R package and database: integration of microRNA–target interactions along with their disease and drug associations Nucleic Acids Research 42, e133–e133

