## Supplementary figures and images for "lncRNA MANCR isoforms selectively mediate multiple levels of epigenomic and P53-responsive transcriptional control in triple negative breast cancer"

### Supplementary Figure 1

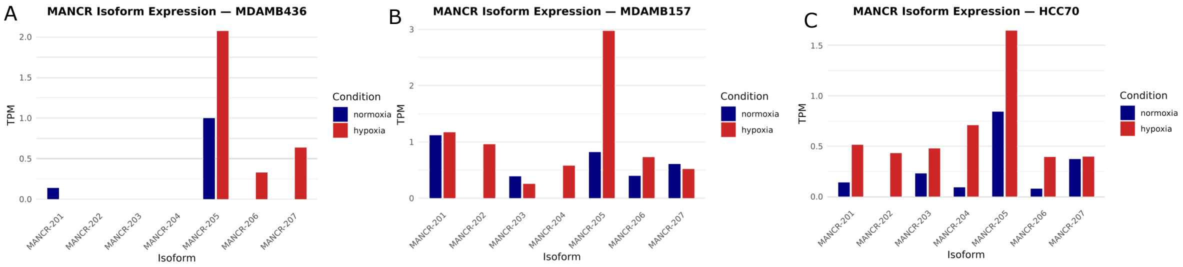

### Supplementary Figure 2

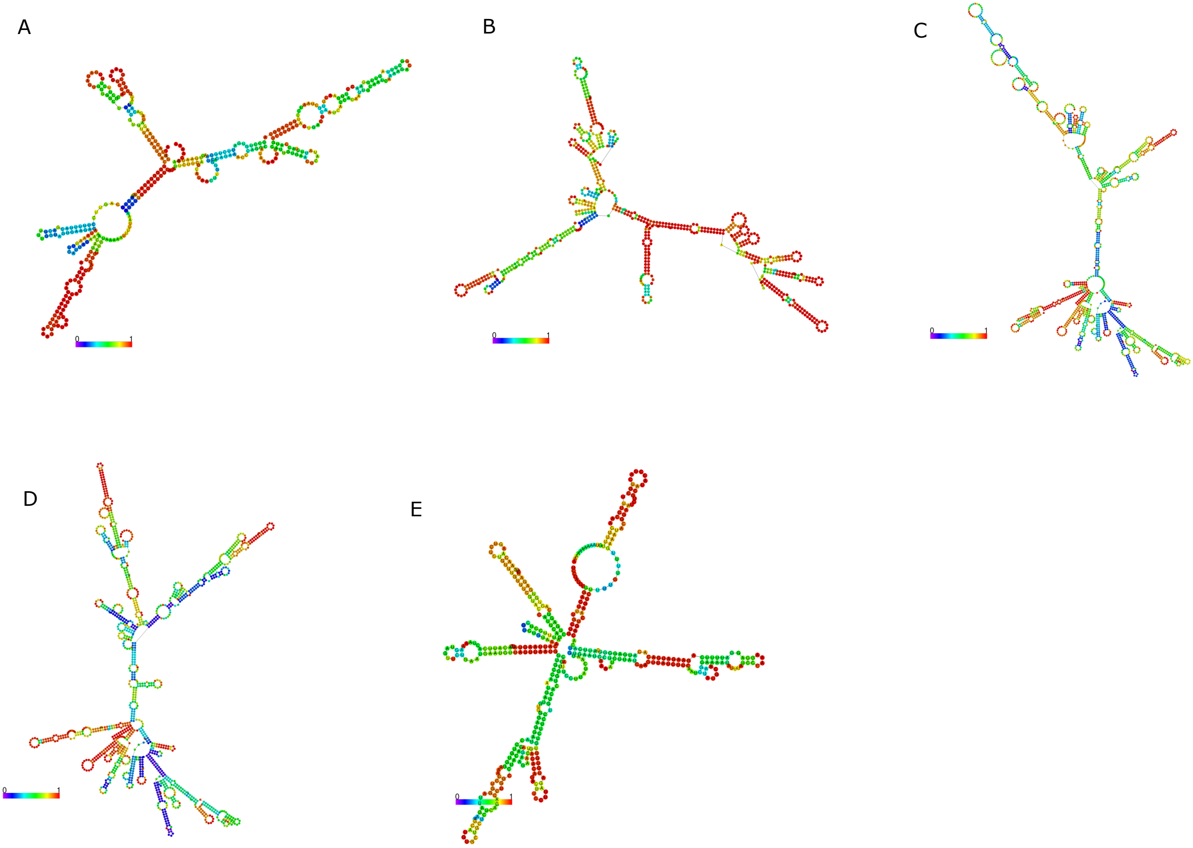

### Supplementary Figure 3

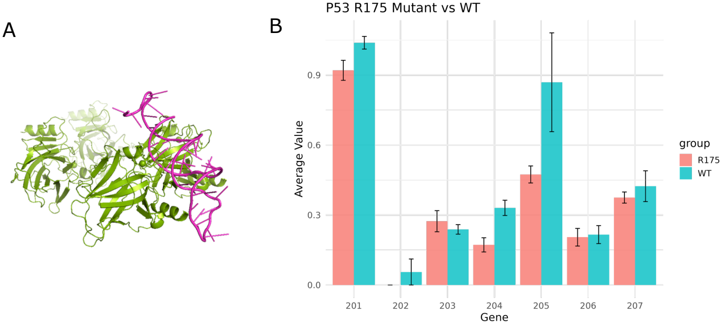

### Supplementary Figure 5

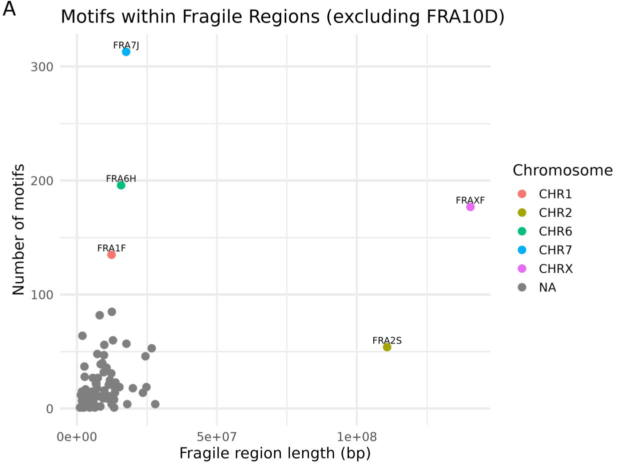

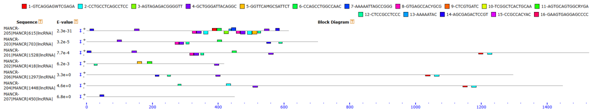

### Supplementary Figure 6

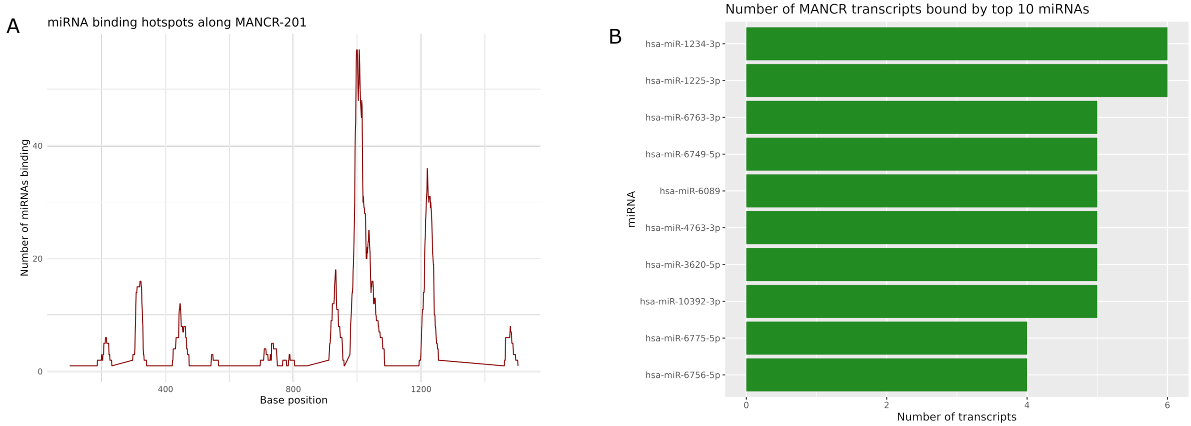
