## Supplementary Figure 4 for "lncRNA MANCR isoforms selectively mediate multiple levels of epigenomic and P53-responsive transcriptional control in triple negative breast cancer"

| Motif | Logo / RC Logo | P-value | E-value | Sites | More | Submit/Download | Positional Distribution / Matches per Sequence |
|---|---|---|---|---|---|---|---|
| 1-GTCAGGAGWTCGAGA | 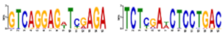 | 1.6e-004 | 2.5e-003 | 340 (27.2%) | ↧ | ⇢ | 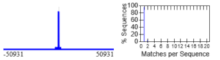 |
| 2-CCTGCCTCAGCCTCC | 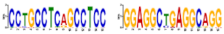 | 2.0e-004 | 3.2e-003 | 441 (35.3%) | ↧ | ⇢ | 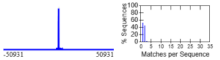 |
| 3-AGTAGAGACGGGGTT | 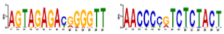 | 2.1e-004 | 3.3e-003 | 359 (28.7%) | ↧ | ⇢ | 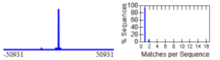 |
| 4-GCTGGGATTACAGGC | 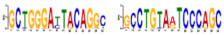 | 3.3e-004 | 5.2e-003 | 420 (33.6%) | ↧ | ⇢ | 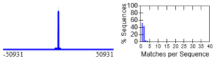 |
| 5-GGTTCAMGCSATTCT | 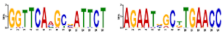 | 1.0e-003 | 1.6e-002 | 269 (21.5%) | ↧ | ⇢ | 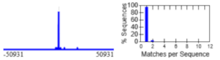 |
| 6-CCAGCCTGGCCAAC | 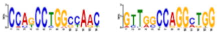 | 1.4e-003 | 2.2e-002 | 467 (37.4%) | ↧ | ⇢ | 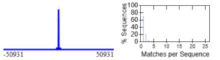 |
| 7-AAAAATTAGCCGGG | 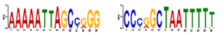 | 4.4e-003 | 7.1e-002 | 306 (24.5%) | ↧ | ⇢ | 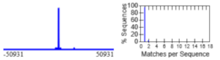 |
| 8-GTGAGCCACYGCG | 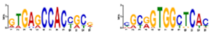 | 4.9e-003 | 7.8e-002 | 255 (20.4%) | ↧ | ⇢ | 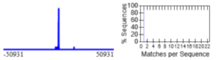 |
| 9-CTCGTGATC | 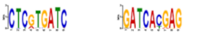 | 6.1e-003 | 9.8e-002 | 190 (15.2%) | ↧ | ⇢ | 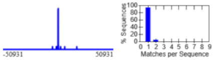 |
| 10-TCGGCTCACTGCAA | 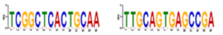 | 7.1e-003 | 1.1e-001 | 292 (23.4%) | ↧ | ⇢ | 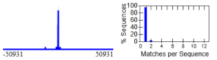 |
| 11-AGTGCAGTGGCRYGA | 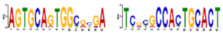 | 1.1e-002 | 1.8e-001 | 196 (15.7%) | ↧ | ⇢ | 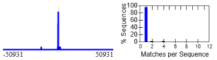 |
| 12-CTCCGCCTCCC | 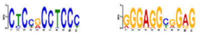 | 2.2e-002 | 3.5e-001 | 369 (29.5%) | ↧ | ⇢ | 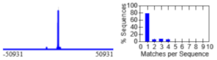 |
| 13-AAAAATAC | 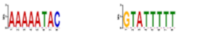 | 3.7e-002 | 6.0e-001 | 259 (20.7%) | ↧ | ⇢ | 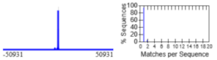 |
| 14-AGCGAGACTCCGT | 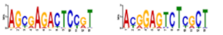 | 8.3e-002 | 1.3e+000 | 174 (13.9%) | ↧ | ⇢ | 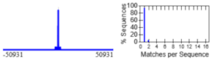 |
| 15-CCGCCACYAC | 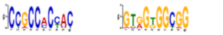 | 9.2e-002 | 1.5e+000 | 274 (21.9%) | ↧ | ⇢ | 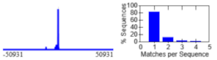 |
| 16-GAAGTGAGGAGCCCC | 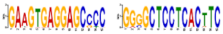 | 4.9e-001 | 7.8e+000 | 103 (8.2%) | ↧ | ⇢ | 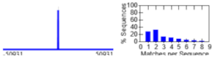 |
